# Ultrasound programmable hydrogen-bonded organic frameworks for sono-chemogenetics

**DOI:** 10.1101/2023.12.08.570721

**Authors:** Wenliang Wang, Yanshu Shi, Wenrui Chai, Kai Wing Kevin Tang, Ilya Pyatnitskiy, Yi Xie, Xiangping Liu, Weilong He, Jinmo Jeong, Ju-Chun Hsieh, Anakaren Romero Lozano, Brinkley Artman, Graeme Henkelman, Banglin Chen, Huiliang Wang

## Abstract

The precise control of mechanochemical activation within deep tissues via non-invasive ultrasound holds profound implications for advancing our understanding of fundamental biomedical sciences and revolutionizing disease treatments. However, a theory-guided mechanoresponsive materials system with well-defined ultrasound activation has yet to be explored. Here we present the concept of using porous hydrogen-bonded organic frameworks (HOFs) as toolkits for focused ultrasound programmably triggered drug activation to control specific cellular events in the deep brain, through on-demand scission of the supramolecular interactions. A theoretical model is developed to visualize the mechanochemical scission and ultrasound mechanics, providing valuable guidelines for the rational design of mechanoresponsive materials at the molecular level to achieve programmable and spatiotemporal activation control. To demonstrate the practicality of this approach, we encapsulate designer drug clozapine N-oxide (CNO) into the optimal HOF nanoparticles for FUS gated release to activate engineered G-protein-coupled receptors in the mice and rat ventral tegmental area (VTA), and hence achieved targeted neural circuits modulation even at depth 9 mm with a latency of seconds. This work demonstrates the capability of ultrasound to precisely control molecular interaction and develops ultrasound programmable HOFs to minimally invasive and spatiotemporally control cellular events, thereby facilitating the establishment of precise molecular therapeutic possibilities. We anticipate that this research could serve as a source of inspiration for precise and non-invasive molecular manipulation techniques, potentially applicable in programming molecular robots to achieve sophisticated control over cellular events in deep tissues.

---

An optimal delivery system should non-invasively and precisely target the specific tissues or cells involved in the disease, achieving a concentration and duration that yield the intended physiological response without excess or inadequate effect.^1,2^ This precision is essential for a wide range of disease treatments, from temporal activation for neural activity modulation,^3,4^ gradual release for treating chronic pain^5^, or the sequential control of drugs at various disease progression stages^6^. Cutting-edge technologies like optogenetics have empowered scientists to execute sophisticated molecular manipulations of opsins at specific cells or projections, significantly advancing our understanding of brain processes and offering the potential for on-demand treatment of neural diseases.^7^ However, their application in deep tissue is constrained due to the inefficient delivery of photons within organisms. Non-invasive molecular manipulation holds great promise for clinical therapeutic applications.

Focused ultrasound (FUS) presents a unique opportunity for non-invasive mechanochemical control in deep tissue with millimetric spatial precision and exemplary safety.^8–12^ Ultrasound-triggered phase-shift microbubbles, nanoemulsions, and sonosensitized liposomes have been developed as promising candidates in mechanotherapy and local anesthesia.^3,13–15^ However, these drug carriers show either insufficient structural stability leading to premature drug release after administration, or excessive stability resulting in an insufficient release of drugs upon ultrasound stimulation.^16^ The development of ultrasound programmable systems with tunable structural stability and ultrasound sensitivity are still challenging. Additionally, the constrained drug loading capacity has also impeded their clinical applicability.^8^ Herrmann *et al* have recently demonstrated the potential of ultrasound to selectively cleave labile covalent or non-covalent bonds, creating new opportunities for precise drug manipulation at the molecular level.^9,17^ However, the presence of strong covalent and non-covalent bonds within polymer frameworks often necessitates high ultrasound power densities, resulting in extended response times on the order of hours.^17–21^ More importantly, the topologically complex nature of these present systems poses challenges in establishing a theoretically predictable system to visualize the intrinsic relationships between scission efficiency, framework molecular structure, and ultrasound power.^22,23^ Despite significant progress in ultrasound-triggered systems, a comprehensive model is yet to be developed to elucidate the manipulation at the molecular level induced by ultrasound (**Supplementary Table 1**). The creation of ultrasound-programmable toolkits is imperative for achieving precise drug activation in optimal clinical disease treatment.

Porous frameworks, including metal-organic frameworks (MOFs) and covalent organic frameworks (COFs), have garnered significant attention as drug delivery platforms due to their remarkable drug loading capacity and well-defined structures.^24,25^ Among them, hydrogen-bonded organic frameworks (HOFs) have recently emerged as a particularly promising class of porous materials with both high structural homogeneity and programmability, self-assembled from organic molecular building units (OMBUs) via hydrogen-bonding and π-π stacking interactions.^26–28^ Unlike strong metal–ligand coordination and covalent bonding interactions in MOFs and COFs, the relatively weak non-covalent interactions make HOFs excellent candidates for mechanochemical activation under FUS stimulation. In addition, the abundant diversities of building units made HOFs easily feasible for fine-tuning their compositions and functionalities to carter custom-design applications.^29^ If the HOFs can be precisely tailored and selectively activated through non-invasive ultrasound, this system could serve as remote medication manipulation, offering precise disease treatment in deep tissue.

In this study, we investigated the potential of HOF nanoparticles to function as mechanoresponsive platforms, enabling ultrasound-programmable and selective activation by adjusting the framework building units (**Fig. 1a**). We developed a theoretical model to elucidate the intrinsic relationship between selective scission and specific ultrasound energy. Through the strategic design of HOF building units, we successfully achieved programmable control over HOF scission for drug activation, enabling activation within varying timeframes, from seconds to hours, depending on the specific application. Additionally, the porous nature of HOFs facilitated high drug loading capacity, reducing the need for carrier materials and minimizing potential adverse effects in medical therapy. We term this approach as UltraHOF, an abbreviation of **ul**trasound **tr**iggered **a**ctivation of **h**ydrogen-bonded **o**rganic **f**rameworks.

**Fig. 1.**
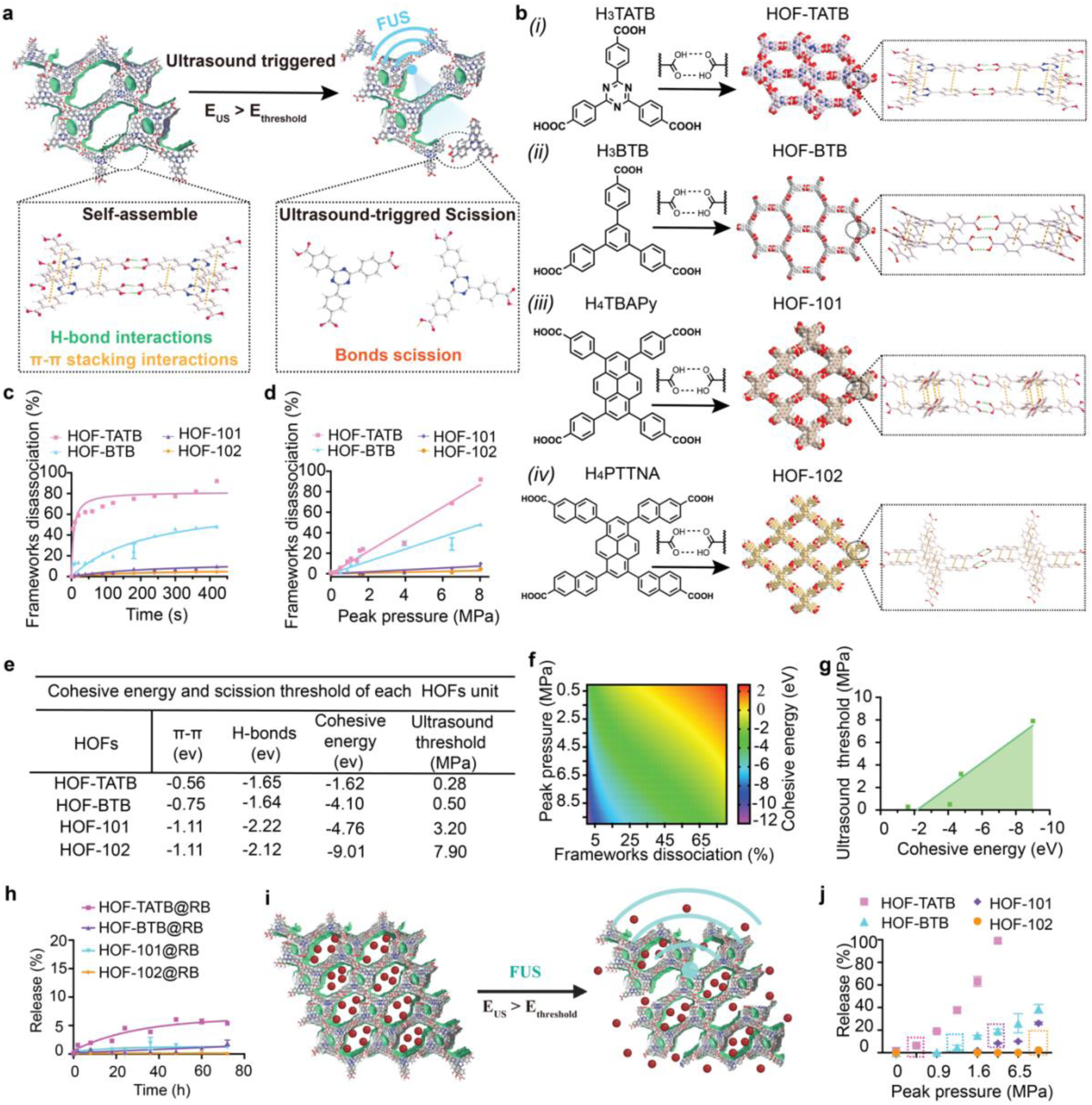
Ultrasound mechanically controlled dissociation of nano-sized hydrogen-bonded organic frameworks (HOFs) in an aqueous solution. (a) Schematic illustration of the ultrasound mechanical stress triggered dissociation of HOFs, where the HOFs were stable in solution but disassociated when triggered by the ultrasound power (E_us_) exceeding HOFs scission threshold (E_us_ > E_threshold_). (b) Four representative organic monomers and the self-assembled porous HOF structures: HOF-TATB, HOF-BTB, HOF-101 and HOF-102. The organic molecular building units (OMBUs) of HOFs self-assemble through hydrogen-bonding and π-π stacking interactions, resulting in the formation of 3D porous frameworks. (c) Ultrasound mechanically triggers the time-resolved dissociation curve of HOFs (mean ± SEM, n=3 independent samples) at peak pressure 8.04 MPa. (d) Ultrasound power dependent dissociation of HOFs (mean ± SEM, n=3 independent samples) at different peak pressure when the equilibrium dissociation of HOFs is achieved. (e) Cohesive energy of different HOFs, including their hydrogen-bonding interactions and π-π stacking interactions in each unit. HOFs were constructed from organic building units through intermolecular non-covalent hydrogen-bonding interactions and π-π stacking interactions (Pi-Pi), where the bonding energy of one unit was denoted as the cohesive energy of HOFs. Moreover, the HOFs displayed varying ultrasound thresholds for disassociation, which were associated with the characteristics of the organic building units. (f) The prediction heatmap of ultrasound-controlled dissociation of HOFs. By referring to the provided heatmap, one can determine the optimal cohesive energy needed to attain a dissociation percentage at a specific ultrasound power. This will guide the structural design of HOFs at the molecular level to achieve the cohesive energy for subsequent programmable control of HOFs dissociation at certain ultrasound peak pressure. (g) Theoretical ultrasound power threshold for triggered dissociation of HOFs. The thresholds for HOFs’ dissociation (HOF-TATB, HOF-BTB, HOF-101, and HOF-102) are 0.28 MPa, 0.50 MPa, 3.20 MPa, and 7.90 MPa. The shadow represents the absence of any dissociation, and the line indicates the theoretical dissociation phase change boundary. (h) The free dye release from HOF nanoparticles without ultrasound stimulation. (i) Schematic diagram of ultrasound triggered drug release from the HOF nanoparticles, with drug release occurs when E_us_ > E_threshold_. (j) Ultrasound triggered dye release from the HOF nanoparticles after 90 s stimulus. The HOF nanoparticles exhibited distinct ultrasound thresholds for drug activation, with the order of sensitivity being HOF-TATB@RB (0.51 MPa) < HOF-BTB@RB (1.55 MPa) < HOF-101@RB (3.94 MPa) < HOF-102@RB (8.04 MPa).

As a proof of concept for this technology, we applied UltraHOF with chemogenetics for *in vitro* and *in vivo* neuromodulation. Specifically, chemogenetics utilized genetically-encoded receptors to activate neurons upon specific drugs agonists binding, exhibiting unique advantages for long period neuromodulation and minimum immunogenicity compared with optogenetics.^30–32^ Recent advancements in remote and minimally invasive control over drug activation offer considerable potential for clinical therapeutic applications in chemogenetics.^8,33,34^ However, the limited temporal control of drug activation and constrained work range in brain tissue still pose significant challenges for precise temporal control of neural activity (**Supplementary Table 2**). As a demonstration of our UltraHOFtechnology, we loaded the CNO into optimal HOF nanoparticles and achieved genetically-targeted neuromodulation through chemogenetics with designer receptors exclusively activated by designer drugs (DREADDs).^30,32,35^ By precisely adjusting the responsiveness of CNO-loaded HOF nanoparticles to FUS in the mice and rat VTA, we effectively attained spatiotemporal control of neural activity even at depth 9 mm with a short latency of mere seconds. Our findings demonstrate that our UltraHOF can achieve high temporal resolution and allow for targeted control of drug activation in specific types of cells while retaining the benefits of minimal invasiveness.

## Characterization of ultrasound scission of mechanoresponsive HOFs

We targeted four different OMBUs with varying density of hydrogen bonding and aromatic rings in respective building units for the synthesis of four porous HOF nanoparticles HOF-TATB, HOF-BTB, HOF-101, and HOF-102 (**Fig. 1b**). The nuclear magnetic resonance (NMR) spectrum of corresponding OMBUs are shown in **Supplementary Fig. 1-4**. We employed a precipitation method to produce nanosized HOFs by fine-tuning solvent polarity. Both dynamic light scattering (DLS) and transmission electron microscopy (TEM) analyses confirmed the successful fabrication of HOF nanoparticles (**Supplementary Fig. 5** and **Supplementary Table 3**). Further powder X-ray diffraction has demonstrated the high crystallinity of nanoscale HOF particles (**Supplementary Fig. 6**). D ue to the planar nature, the OMBUs spontaneously assembled into two-dimensional layers via carboxylic acid H-bonded dimers, where the tetracarboxylic OMBUs forms a square lattice topology layer in HOF-101 and HOF-102, while the tricarboxylic ones crystalize into hexagonal honeycomb layers in HOF-TATB and HOF-BTB. More specifically, the strong π-π stacking of pyrene moieties in HOF-101 and HOF-102 drives the 2D square layers to further stack with adjacent layers into 3D crystals with rhombus 1D channel (**Fig. 1 and Supplementary Table 4**).^36,37^ In contrast to the tetracarboxylic building units, the stacking of the self-assembled tricarboxylic acids in HOF-BTB and HOF-TATB is inherently weaker. Therefore, the layers are first stacked to align the aromatic motifs and further interwoven to maximize the overall molecular packing.^29,38^ The experimental 77 K nitrogen adsorption tests determined that HOF nanoparticles exhibit characteristic porosity (**Supplementary Fig. 7**).

We next verified the ultrasound-triggered scission of HOF nanoparticles in an aqueous medium by monitoring the framework dissociation percentage over time (**Fig. 1c** and **Supplementary Fig. 8**). Irradiation of the HOF nanoparticles with ultrasound (1.5 MHz) at a power of 8.04 MPa led to approximately 91.8%, 45.3%, 8.4%, and 4.7% dissociation after equilibrium in HOF-TATB, HOF-BTB, HOF-101, and HOF-102, respectively. However, no obvious dissociation was observed even when these HOFs were heated at 100 °C for 5 minutes, demonstrating their high thermal stability (**Supplementary Fig. 9**). These results determined that the ultrasound stress, rather than thermal effect, constitutes the main driving force to shear the intramolecular noncovalent bonds and dissociate the frameworks. In fact, HOFs demonstrate an excellent capability for ultrasound power programmable behavior through mechanochemical activation. Activation of HOFs occurs exclusively when the ultrasound power reaches a specific peak pressure, with the ultrasound power activation thresholds being contingent upon the type of OMBU used (**Supplementary Fig. 8**). In addition, the dissociation percentage increases in response to higher ultrasound peak pressure for all 4 different HOFs, with difference in increasing rate depending on the specific type of HOF, as shown in **Fig. 1d**.

The structure-property relationships of the HOFs were further investigated using density functional theory (DFT) calculations, by calculating the cohesive energy (*E_cohesive_*, the energy needed to dissociate the HOF crystals to isolated building units) of, from the lowest to the highest, HOF-TATB, HOF-BTB, HOF-101, and HOF-102 in solution, indicating increasing relative stability of the HOFs (**Fig. 1e**). This observation is consistent with the experimental findings where the dissociation equilibrium constant (*k,* the ratio of dissociated HOF building units percentage to undissociated HOFs percentage at certain ultrasound pressure) decreases sequentially in these four HOFs (**Supplementary Table 5**). Our reaction model indicated a linear relationship between ln(*k*) and *E_cohesive_* of the HOFs (**Supplementary Fig. 10**), and thus we can extrapolate the required minimum *E_cohesive_* of HOF at any desired ln(*k*). Additionally, we found a linear relationship between the required minimum *E_cohesive_* and the ultrasound peak pressure (*E_US_*) of the HOFs (**Supplementary Fig. 11**, **Supplementary Table 6**) at any ln(*k*). From these linear relationships between ln(*k*) and *E_cohesive_*, and between *E_US_* and *E_cohesive_*, we developed a three-variable model to represent how these variables are correlated. The heat map in **Fig. 1f**, predicts the *E_cohesive_* of HOFs needed for achieving a given dissociation percentage at a specific ultrasound power, thus providing a guideline for designing HOFs for on-demand drug activation by ultrasound. From the heat map, we also determined the theoretical ultrasound power thresholds (the minimum ultrasound peak pressure to achieve 3% dissociation of HOFs) for HOFs (HOF-TATB, HOF-BTB, HOF-101, and HOF-102) are 0.28 MPa, 0.50 MPa, 3.20 MPa, and 7.90 MPa respectively (**Fig. 1e,g**). Since the sensitivity of HOFs to ultrasound is governed by the overall strength of weak interactions encoded in the HOF structure, primarily contributed by hydrogen-bonds and π-π interactions among OMBUs, the relationship between the OMBU structures and ultrasound sensitivity is important for elucidating the acoustic scission principles of HOFs at the molecular level. The DFT results (**Fig. 1e**) show that the hydrogen-bonding energy (*E_HB_*, the energy per hydrogen-bonded dimer) is approximately −1.1 eV and remains relatively stable across the molecular structures of the OMBU in the four HOFs. However, the number of hydrogen bonds in each OMBU varies with the carboxyl groups. Specifically, each OMBU in HOF-TATB and HOF-BTB form 1.5 H-bonds, while those of HOF-101 and HOF-102 form 2 H-bonds each. This difference contributes to the higher hydrogen-bond energy in HOF-101 and HOF-102 compared to HOF-TATB and HOF-BTB. Importantly, the increased number of hydrogen bonds in each OMBU significantly enhances the ultrasound-triggered stability of HOF nanoparticles. For π-π interactions, the energy (*E*_π-π_) varies with the structure of the OMBU. Higher *E*_π-π_ was found in HOF-101 and HOF-102 due to the increased number of aromatic fused rings in the OMBU (**Fig. 1e**). Since mechanochemical bond scission is more likely to occur in weak bonds to trigger the dissociation of frameworks, it is reasonable to suggest that the OMBUs with fewer H-bonds and aromatic fused rings in the backbone should be used for HOFs with a higher ultrasound sensitivity.

## Ultrasound programable drug activation and *in vitro* neuromodulation

These HOF nanoparticles exhibited outstanding drug loading capacity. As shown in **Supplementary Table 7**, the loading capacity increased in HOF-TATB (15.1 ± 1.4%), HOF-BTB (15.8 ± 2.7%), HOF-101 (27.0 ± 1.5%), and HOF-102 (29.8 ± 1.3%) due to their increased porosity.^29,36–38^ This characteristic reduces the drug carriers needed in delivering specific drug concentrations, thereby minimizing the side effects. We next utilized dye release experiments to examine the free drug release of the HOF nanoparticles without ultrasound application (**Fig. 1h**). Specifically, only 5.5 ± 0.1% of the dye was prematurely released from HOF-TATB nanoparticles without ultrasound even after 3 days of incubation, and the premature release percentage of dye could be reduced further to 1.9 ± 0.9%, 1.2 ± 0.2%, 0.1 ± 0.1% in HOF-BTB, HOF-101 and HOF-102 due to their increased *E_cohesive_*. In addition, ultrasound-triggered release experiments demonstrated that the percentage of drug released increased with the ultrasound peak pressure used and the type of HOF with lower *E_cohesive,_* (**Fig. 1i,j** and **Supplementary Fig. 12**), consistent with the theoretical ultrasound power thresholds conducted from our prediction heat map (**Fig. 1f**). Of note, among various HOF nanoparticles, HOF-TATB nanoparticles exhibited the highest sensitivity to ultrasound, providing optimal temporal resolution for drug activation gating with minimum ultrasound pressure while HOF-102 nanoparticles exhibited the highest stability and drug loading capacity (**Supplementary** Fig.12 and **Supplementary Table 7**). For demonstration of the UltraHOF for deep brain stimulation with high temporal resolution, we decided HOF-TATB to minimize the ultrasound pressure needed.

After drug loading, HOF-TATB nanoparticles did not show any notable alterations in size and morphology (**Fig. 2a**). Additionally, their negative surface potential ensured excellent biostability, even in the presence of 10% fetal bovine serum (**Supplementary Fig. 13**). However, it is worth noting that as the size decreased, there were morphological changes of HOF nanoparticles after exposure to ultrasound stimulation (**Fig. 2a** and **Supplementary Table 3**). We observed drug release was triggered under the ultrasound stimulation within clinical safe range (1.5 MHz, 1.5 MPa), whereas no significant release occurred without ultrasound (**Fig. 2b and Supplementary Fig. 14**). The percentage of released drugs also increases with the ultrasound peak pressure, with an activation threshold of around 0.51 MPa (**Fig. 2c and Supplementary Fig. 14**). Moreover, dye release could be repeatedly triggered from the HOF-TATB nanoparticles by a repeated stimulus (10-second pulse), resulting in an example of four triggerable events releasing 10.5 ± 1.6%, 5.0 ± 3.0%, 6.5 ± 5.2% and 10.0 ± 4.4% respectively (**Fig. 2d**). We also determine that the payload release percentage increases linearly with the frameworks dissociation percentage (**Supplementary Fig. 15**). Furthermore, we determined the biosafety of the HOF-TATB nanoparticles through the hemolysis and cell viability tests. The results revealed no apparent toxicity or hemolysis, even at high concentrations. This suggested that our HOF-TATB nanoparticles are generally biocompatible and biosafe as drug delivery platforms (**Supplementary Fig. 16**).

**Fig. 2.**
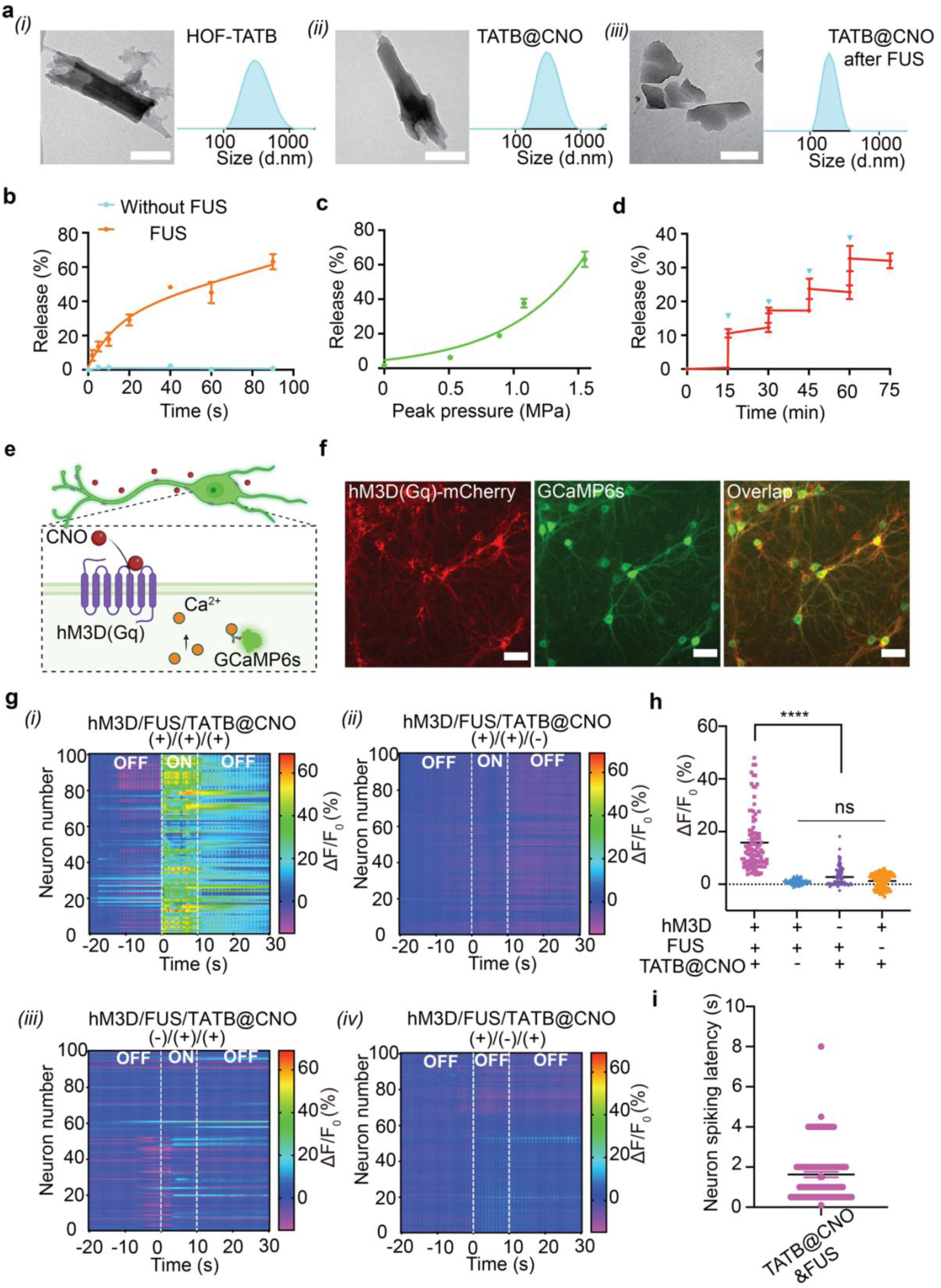
Ultrasound controlled cargo release from HOF-TATB nanoparticles and their *in vitro* modulation of neural activity. (a) TEM images and hydrodynamic size distribution measured by dynamic light scattering of the HOF-TATB nanoparticles (i) before loading of clozapine-N-oxide (CNO) and (ii) after loading of CNO, and (iii) after irradiating by ultrasound (1.5 MHz, 1.55 MPa, 60 s). Scale bar: 200 nm. (b) Ultrasound triggered dye release (mean ± SEM, n=3 independent samples) from HOF-TATB at 1.5 MHz, 1.55 MPa. (c) Ultrasound triggered dye release (mean ± SEM, n=3 independent samples) from HOF-TATB after 60 s irradiation (1.5 MHz) at different peak pressures. (d) Repeated ultrasound triggered drug release. The blue arrows indicate ultrsound stimulus (1.5 MHz, 1.55 MPa, pulse 10 s). Mean ± SEM; more than 3 times independent tests (n ≥ 3). (e) Ultrasound triggered CNO release from the HOF-TATB nanoparticles for hM3D(Gq) expressing neuron activation. (f) Fluorescence images of the primary cortical neurons expressing hSyn::hM3D (Gq)-mCherry and hSyn::GCaMP6s-WPRE-SV40. Scale bar: 40 μm. (g) Heat maps of normalized GCaMP6s fluorescence intensity from 100 neurons in different experimental conditions, including (i) hM3D(+)/FUS(+)/TATB@CNO(+), (ii) hM3D(+)/FUS(+)/TATB@CNO(-), (iii) hM3D(-)/FUS(+)/TATB@CNO(+) and (iv) hM3D(+)/FUS(-)/TATB@CNO(+). The experiments were independently repeated at least three times. F_0_ is the mean fluorescence intensity of baseline, hM3D(Gq)^+^ means neurons with hM3D(Gq) expression, hM3D(Gq)^-^ means neurons without hM3D(Gq) expression, FUS^+^ means the neurons are treated with the focused ultrasound and FUS^-^ means the neurons without focused ultrasound treatment, TATB@CNO^+^ means neurons with CNO loaded HOF-TATB nanoparticles, and TATB@CNO^-^ means neurons HOF-TATB nanoparticles only without CNO. OFF means ultrasound is off. ON means ultrasound is on (1.5 MHz, 1.08 MPa, pulse 10 s). (h) Statistical analysis of calcium signal changes in 100 primary neurons under the different conditions. Mean ± SEM, n ≥ 3 independent tests. Two-way ANOVA and Tukey’s multiple comparison test (P ≥ 0.05 (ns), * 0.01≤ P < 0.05, ** 0.001≤ P < 0.01, **** P< 0.0001). (i) Normalized *in vitro* neuron spiking latency under sono-chemogenetics stimulation. The hM3D(Gq) and GCaMP6s expressing neurons (100 neurons) were conducted to analyze the latency between the neuron firing and ultrasound stimulation. The results determined that the latency of sono-chemogenetics stimulation is around 1.6 s. Mean ± SEM; 3 times independent tests (n ≥ 3).

Next, we applied our HOF-TATB nanoparticles for loading of CNO (TATB@CNO) for ultrasound-triggering of CNO to activate DREADDS in cultured neurons (**Fig 2e**). First, the neurons were transduced with endogenous designer receptors hM3D(Gq) with red fluorescent reporter mCherry (AAV9-hSyn-hM3D(Gq)-mCherry) and green fluorescence calcium indicator (AAV9-hSyn-GCaMP6s-WPRE-SV40), as shown in **Fig 2f**. We initially validated the activation of hM3D(Gq) expressing neurons with the free designer drug CNO using fluorescence imaging of GCaMP6s calcium indicators (**Supplementary Fig. 17**). Then, by subjecting CNO-encapsulating nanoparticles (TATB@CNO) to ultrasound irradiation, over 90% of hM3D(Gq)^+^ neurons fired with a latency of 1.6 s and continuously activated for ∼60 s. However, only sporadic neuronal activation was observed in the absence of ultrasound stimulus, hM3D(Gq) expression or TATB@CNO nanoparticles (**Fig. 2g-i** and **Supplementary Fig. 18**). The burst release of CNO from HOF-TATB nanoparticles under ultrasound stimulus enables the rapid triggering of hM3D(Gq), inducing long-term (> 60 s) neuron membrane depolarization. Our UltraHOF-enabled sono-chemogenetics provides a novel method for achieving fast and continuous neuronal activation with minimal invasiveness.^7^

## UltraHOF-enabled sono-chemogenetic deep brain stimulation in mice

Precisely timed activation of genetically targeted neurons is important for researchers to understand the links between brain activity and behavior.^7,39^ We subsequently assessed real-time UltraHOF-enabled sono-chemogenetic neural excitation through fiber photometry in the ventral tegmental area (VTA) of mice, a region known for its role in regulating reward learning and depression.^40^ Ultrasound energy propagates through tissue as a traveling pressure wave, with penetration depth increasing with lower frequency but at the cost of decreased resolution.^41^ In our experiments, a 1.5 MHz transducer achieved a maximum penetration depth of 20 mm, with 37% delivery efficiency at a tissue depth of 10 mm (**Supplementary Fig. 19**). The ultrasound energy heatmap in the mouse brain showed that a primary power of 1.4 MPa produced an acoustic pressure of around 0.9 MPa at the VTA of mice (**Supplementary Fig. 20**), which is sufficient to activate CNO release (**Fig. 2c** and **Supplementary Fig. 14**). We performed unilateral transduction of neurons in VTA using the AAV9-hSyn-hM3D(Gq)-mCherry and AAV9-hSyn-GCaMP6s-WPRE-SV40 virus, followed by the injection of TATB@CNO in the same region 4 weeks later (**Fig. 3a,b**). We used fiber photometry to record Ca^2+^ influx-induced GCaMP6s signal increase in mice to assess neuronal excitation in response to ultrasound-gated release of CNO after 2 days of injection. The GCaMP6s signal increased significantly upon ultrasound irradiation of the VTA region for neurons with hM3D(Gq) expression, but remained unchanged without hM3D(Gq) expression, TATB@CNO nanoparticles or ultrasound stimulus (**Fig. 3c,d**). Our sono-chemogenetics approach demonstrated high temporal resolution, with a 3.5 s latency for neural activations upon ultrasound stimulus (**Fig. 3e**). We observed over 120 s of continuous excitation of neural activity with each only 10 s sonochemogenetic stimulation, and multiple increases in GCaMP6s fluorescent signal were observed upon repeated exposure to ultrasound even after 5 days injection due to the long-term biostability of TATB@CNO in the brain (**Supplementary Fig. 21, 22**). Immunofluorescence analysis of the expression of c-fos, an immediate early gene and a marker for neural excitation,^42^ further revealed that neurons excitation in the VTA was only triggered by the FUS stimulus in mice with both expression of hM3D(Gq) and injected TATB@CNO nanoparticles (**Fig. 3f-g**).

**Fig. 3.**
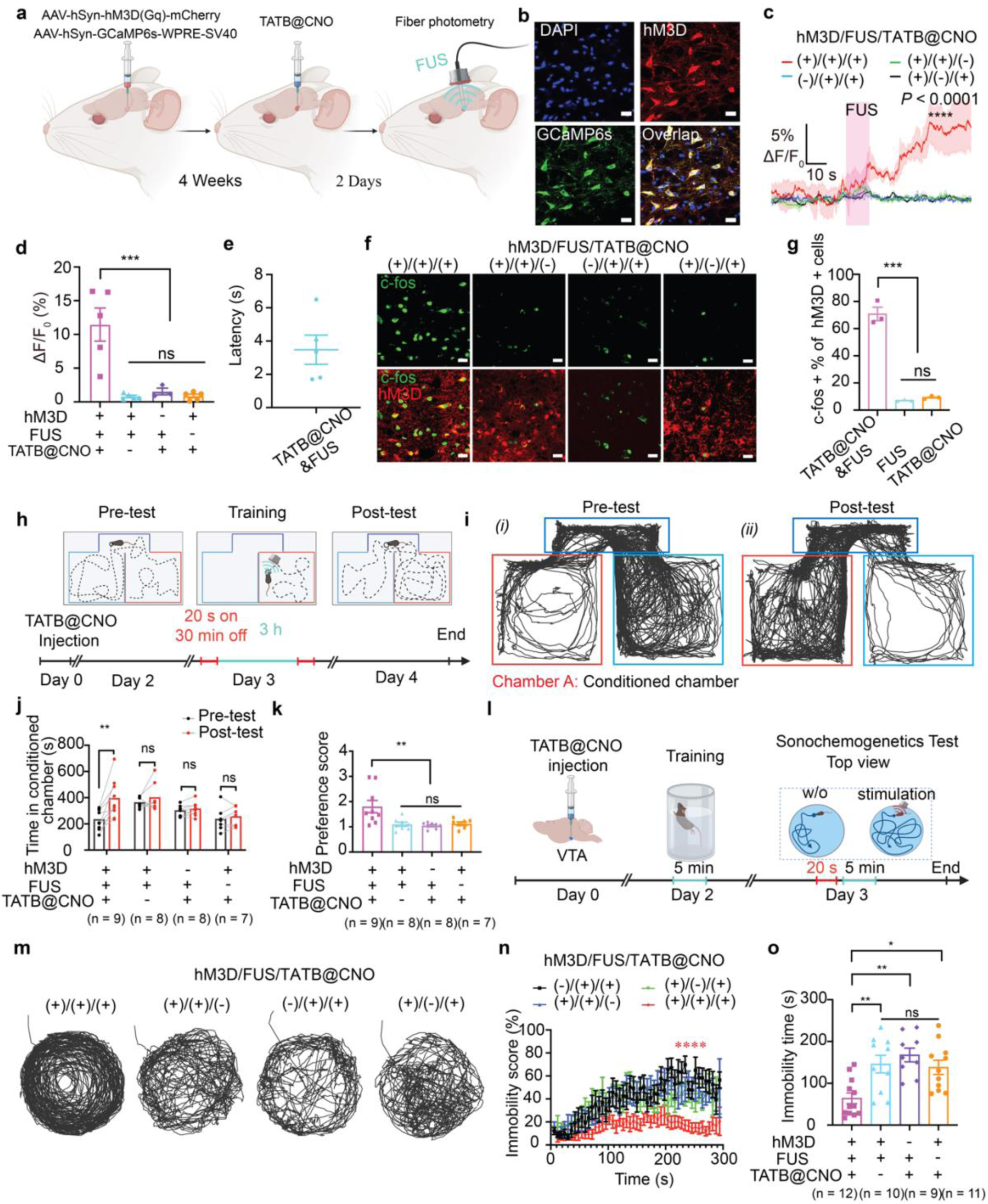
*In vivo* sono-chemogenetic deep brain stimulation in mice. (a) Experimental scheme of the *in vivo* fiber photometry in the VTA area. (b) Confocal fluorescence images of the co-expression of hM3D(Gq) and GCaMP6s in mouse VTA area. Scale bar: 20 μm. (c) Normalized GCaMP6s fluorescence intensity change (ΔF/F_0_) in mice VTA under the different experimental conditions. Remarkable fluorescence increase was observed only when the FUS was applied to the mice with hM3D(Gq) expressed in VTA neurons. The pink area represents the FUS irradiation (1.5 MHz, 1.40 MPa, pulse 10s). Solid line, mean; shade area, SEM; at least 3 mice in each group (n ≥ 3). One-way ANOVA and Tukey’s multiple comparison test (P ≥ 0.05 (ns), * 0.01≤ P < 0.05, ** 0.001≤ P < 0.01, **** P< 0.0001). (d) Statistical analysis of calcium signal changes in mice VTA region under different conditions. Mean ± SEM, n ≥ 3 independent mouse. Two-way ANOVA and Tukey’s multiple comparison test (P ≥ 0.05 (ns), * 0.01≤ P < 0.05, ** 0.001≤ P < 0.01, **** P< 0.0001). (e) Normalized *in vivo* neuron spiking latency in mice VTA under sono-chemogenetics stimulation, where the sono-chemogenetics exhibited temporal resolution with around 3.5 s latency. (f) c-fos expression confocal fluorescence images in the VTA after the mice are treated with different conditions. Remarkable increase of c-fos expression was observed only when the mice with hM3D(Gq) expression in VTA neurons were treated with FUS (1.5 MHz, 1.40 MPa, pulse 20 s, focus length 5 mm). Scale bar: 20 μm. (g) Quantification of the c-fos expression percentage among hM3D(Gq)+ neurons. Mean ± SEM, n ≥ 3 independent samples. Two-way ANOVA and Tukey’s multiple comparison test (P ≥ 0.05 (ns), * 0.01≤ P < 0.05, ** 0.001≤ P < 0.01, **** P< 0.0001). (h) Scheme of conditioned place preference tests with sono-chemogenetics. (i) Traces of mouse freely exploring apparatus (*i*) before and (*ii*) after sono-chemogenetic stimulation. Red one is the conditioned chamber. (j) Statistical analysis of time spent in the FUS stimulation chamber. Mean ± SEM, n, number of the mice in each group. Paired t tests and two-tailed comparison test (P ≥ 0.05 (ns), * 0.01≤ P < 0.05, ** 0.001≤ P < 0.01, **** P< 0.0001). (k) preference score of mice with different conditions. Mean ± SEM, n, number of the mice in each group. Two-way ANOVA and Tukey’s multiple comparison test (P ≥ 0.05 (ns), * 0.01≤ P < 0.05, ** 0.001≤ P < 0.01, **** P< 0.0001). (l) Scheme of forced swimming test (FST) with sono-chemogenetics. (m) Presentative traces of mice in FST under the different conditions. (n) Time-resolved mouse immobility score curve and (o) statistical analysis of immobility time of mice in the FST tests. Only the mice with hM3D(Gq) expression in VTA neurons and TATB@CNO nanoparticles only exhibited reduced immobility score after FUS stimulation. Mean ± SEM, n, number of the mice in each group. Two-way ANOVA and Tukey’s multiple comparison test (P ≥ 0.05 (ns), * 0.01≤ P < 0.05, ** 0.001≤ P < 0.01, **** P< 0.0001).

Based on the notable neural excitation observed in the VTA, we subsequently assessed the ability of UltraHOF-enabled sono-chemogenetics in modulating the reward learning behavior of mice (**Fig. 3h**). After injecting TATB@CNO nanoparticles in the VTA region with hM3D(Gq) expression on Day 0, mice were allowed to freely explore the apparatus without any stimulus on Day 2 (Pre-tests). On Day 3, mice received sonochemogenetic stimulus (1.5 MHz, pulse 20 s, 1.40 MPa) in the designated conditioned chambers with two sessions. On Day 4, mice again freely explored the apparatus without any stimulus (Post-test). Trail tracing revealed that mice expressing hM3D(Gq) preferentially explored the conditioned side of the apparatus after receiving sono-chemogenetics (**Fig. 3i**), resulting in an approximately 2-fold increase in time spent in the conditioned chamber (**Fig. 3j**). Notably, in the absence of hM3D(Gq) expression, TATB@CNO nanoparticles, or ultrasound stimulus, the preference did not vary significantly within the experimental trial (**Fig. 3j,k**).

Based on the previous evidence of the antidepressant effects of neural activity in the VTA,^40^ we aimed to investigate whether UltraHOF-enabled sono-chemogenetics could impact mouse behavior in the forced swim test (FST). To achieve this, VTA neurons were transduced with hM3D(Gq) and mice were allowed to recover from TATB@CNO VTA injection surgery (Day 0) for two days before undergoing a 5-minute pre-FST test (Day 2). On Day 3, mice received ultrasound stimulus (1.5 MHz, pulse 20 s, 1.40 MPa) before the 5-minute FST assay (**Fig. 3l**). We tracked the mice’s swimming patterns during the FST period using motion cameras. The results showed that hM3D(Gq)+ mice exhibited increased activity, increased swimming distance during the FST period (**Fig. 3m**). Furthermore, the dynamic motion analysis showed that hM3D(Gq)+ mice exhibited higher mobility throughout the FST period, with a slight increase in immobility preference only after swimming for 2 minutes. On the other hand, in the absence of hM3D(Gq) expression, TATB@CNO nanoparticles, or ultrasound stimulus, the immobility time significantly increased in mice (**Fig. 3n-o**). These findings suggest that UltraHOF-enabled sono-chemogenetics can effectively modulate mouse behavior in the FST. After 14 days of stimulation, we also evaluated the *in vivo* biocompatibility of our sono-chemogenetics approach by immunostaining brain sections. These results demonstrate that this approach did not cause any significant cell toxicity through H&E staining (**Supplementary Fig. 23**). Additionally, we did not observe any activation of microglia or astrocytes, nor did we detect any neuron apoptosis (**Supplementary Fig. 24**-26).

## UltraHOF-enabled sono-chemogenetic deep brain stimulation in rats

While mice are smaller, cheaper and available with more transgenic types for neuroscience research, rats are paramount to clinical translation because of their thicker skulls and larger depths of brain tissue that more closely meet practical application requirements in clinical. Minimally-invasive, genetically-targeted brain modulation has been successfully demonstrated in mice using magnetic nanotransducers,^4,34,43^ sonogenetics,^44,45^ X-ray activated systems,^46^ and NIR-based approaches.^47,48^ However, it is still a challenge to achieve genetically targeted deep brain neuromodulation and behavior control in rats. Utilizing the high energy transmission efficiency of ultrasound in tissues and the high ultrasound sensitivity of HOF-TATB nanoparticles, we investigated the efficacy of UltraHOF-enabled sono-chemogenetics in rats. Our analysis of the rat head ultrasound heatmap revealed that a peak ultrasound pressure of 1.30 MPa was achieved to gate the CNO release, by applying a primary ultrasound power of 2.45 MPa (**Supplementary Fig. 27**). Prior to TATB@CNO nanoparticle injection and optical fiber implantation, hM3D(Gq) and GCaMP6s were transduced into the rat VTA neurons (**Fig. 4a,b**). After a recovery period of two days, we recorded the green fluorescent signal of GCaMP6s under ultrasound stimulation (1.5 MHz, pulse 20 s, 2.45 MPa) to evaluate the change in neuron activity. We observed a significant increase in GCaMP6s signal in hM3D(Gq)+ rats with stimulation, but not in the absence of hM3D(Gq), TATB@CNO nanoparticles, or ultrasound stimulus (**Fig. 4c,d**). Notably, our UltraHOF-enabled sono-chemogenetics achieved neuron activation with a latency of 8.8 s from the application of ultrasound and continuous activation for over 60 s (**Fig. 4e**). In addition, we evaluated neuron activation in post-hoc brain slices via c-fos immunostaining. Remarkable c-fos signals were observed in neurons expressing hM3D(Gq) in VTA, whereas few c-fos signals were observed in the absence of hM3D(Gq), TATB@CNO nanoparticles, or ultrasound stimulus (**Fig. 4f-g**). These results confirm that our approach can achieve remote brain stimulation even in deep brain regions of rats.

**Fig. 4.**
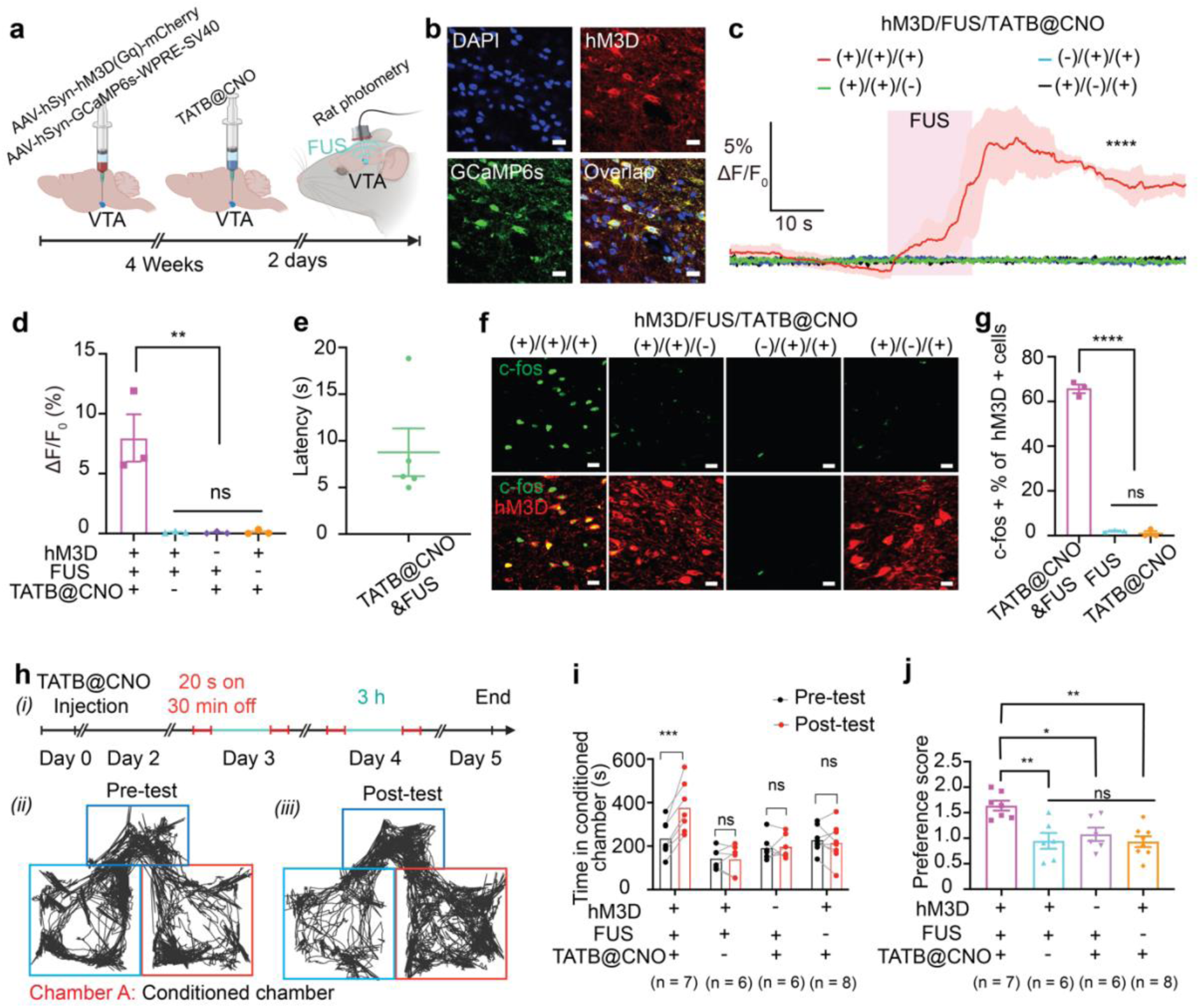
*In vivo* sono-chemogenetic deep brain stimulation in rats. (a) Experimental scheme of the *in vivo* fiber photometry in rat VTA area. (b) Confocal fluorescence images of the co-expression of hM3D(Gq) and GCaMP6s in rat VTA area. Scale bar: 20 μm. (c) Normalized GCaMP6s fluorescence intensity change (ΔF/F_0_) in rat VTA under the different experiment conditions. Remarkable fluorescence increase was observed only when the FUS was applied to treat the mice with hM3D(Gq) expressed in VTA neurons. The pink area represents the FUS irradiation ((1.5 MHz, 2.45 MPa, pulse 20 s, focus length 9 mm). Solid line, mean; shade area, SEM; at least 3 rats in each group (n ≥ 3). One-way ANOVA and Tukey’s multiple comparison test (P ≥ 0.05 (ns), * 0.01≤ P < 0.05, ** 0.001≤ P < 0.01, **** P< 0.0001). (d) Statistical analysis of calcium signal changes in rat VTA region under the different conditions. Mean ± SEM, n ≥ 3 independent rats. Two-way ANOVA and Tukey’s multiple comparison test (P ≥ 0.05 (ns), * 0.01≤ P < 0.05, ** 0.001≤ P < 0.01, **** P< 0.0001). (e) Normalized *in vivo* neuron spiking latency in rat VTA under sono-chemogenetics stimulation, where the sono-chemogenetic stimulation exhibited around 8.8 s latency at depth of 9 mm. (f) c-fos expression in confocal fluorescence images in the VTA after the rat is treated with different conditions. Scale bar: 20 μm. (g) Quantification of the c-fos expression percentage among the hM3D(Gq)+ neurons. Mean ± SEM, n ≥ 3 independent samples. Two-way ANOVA and Tukey’s multiple comparison test (P ≥ 0.05 (ns), * 0.01≤ P < 0.05, ** 0.001≤ P < 0.01, **** P< 0.0001). (h) Rat conditioned place preference tests, (*i*) scheme of conditioned place preference tests with sono-chemogenetics. Traces of mouse freely exploring apparatus (*ii*) before and (*iii*) after sono-chemogenetic stimulation. Red one is the conditioned chamber. (i) Statistical analysis of time spent in the FUS stimulation chamber. Mean ± SEM, n, number of the rat in each group. Paired t tests and two-tailed comparison test (P ≥ 0.05 (ns), * 0.01≤ P < 0.05, ** 0.001≤ P < 0.01, **** P< 0.0001). (j) preference score of rats at different conditions. Mean ± SEM, n, number of the rats in each group. Two-way ANOVA and Tukey’s multiple comparison test (P ≥ 0.05 (ns), * 0.01≤ P < 0.05, ** 0.001≤ P < 0.01, **** P< 0.0001).

Lastly, we further assessed the potential of UltraHOF-enabled sono-chemogenetics in shaping behaviors in rats using a 4-day conditioned place preference assay. Rats transduced with hM3D(Gq) in the VTA region received TATB@CNO nanoparticles on Day 0 and were allowed to freely explore the chambers after a two-day recovery period. On Day 3 and Day 4, the rats were subjected to ultrasound stimulus (1.5 MHz, pulse 20 s, 2.45 MPa) in the designated conditioned chambers, with two sessions each day. On Day 5, post-tests were conducted, where the rats were allowed to freely explore the chamber without any stimulus (**Fig. 4h**). The rats treated with our sono-chemogenetics exhibited a preference to stay in the conditioned chamber compared to other groups in post-tests (**Fig. 4i**). Our results demonstrate a statistically significant difference in the preference score between the hM3D+/FUS+/TATB@CNO+ group and the other control groups without hM3D(Gq), TATB@CNO nanoparticles, or ultrasound stimulus (**Fig. 4j**), indicating that our sono-chemogenetics can effectively modulate reward learning behaviors in rats through non-invasive sono-optogenetic deep brain stimulation.

## Conclusions

This work presents an ultrasound-activated HOF nanomaterial system with finely tuned interactions at the molecular level through modifying the chemical structure of interaction units. Specifically, HOFs hold together via weak intermolecular hydrogen bond and π-π interactions between each discrete organic molecule units to form 3D porous frameworks, hence giving them tunable stability in aqueous conditions, high loading capacity and ultrasound programmable disassociation. Ultrasound stress provided the main driving force to programmably shear the intramolecular noncovalent bonds to achieve controlled mechanochemical activation. Through the manipulation of hydrogen bond density and the number of aromatic fused rings in the backbone structures of the organic ligands, a theoretical model is developed to elucidate the structure and functionality relationships in the HOFs, providing valuable guidelines for the precise and rational design of HOF building units at the molecular level to achieve on-demand and programmable drug activation under a desirable ultrasound pressure.

Given such abilities of HOFs, ultrasound-triggered temporal and programmable drug activation opened a new realm of non-invasive neural control and medical therapy, such as chemogenetic modulation of targeted neural circuits demonstrated in this study. By tuning HOFs nanoparticles sensitivity to respond to focused ultrasound, we successfully achieve spatiotemporal control of deep brain neural circuits in both mice and rats with a latency of only seconds. The results demonstrate that UltraHOF-enabled sono-chemogenetics can achieve a high temporal resolution, long-period neuromodulation while retaining the benefits of minimal invasiveness. Our findings have demonstrated that our UltraHOF technology has the combination of high drug loading content, high biostability, low immunogencity and unique ultrasound programmability for non-invasive, precise medication therapy. In addition to its application for sono-chemogenetics, the UltraHOF technology are capable of releasing different types of molecules with designable medication activation sensitivity and resolution. This enables precise and non-invasive control of various cellular events in deep tissues.

## Methods

### HOF nanoparticles preparation

#### a. HOF-TATB nanoparticles

30 mg H_3_TATB was dissolved in 3 mL dimethylformamide (DMF), and then 12 mL distilled water was added with a stirring speed of 1000 rpm. After stirring for 10 min, the nanoparticles were collected via centrifugation at 12000 rpm (13523 x g) for 5 min (Centrifuge 5420, Eppendorf), and then washed with acetone and distilled water 3 times, respectively. The final products with around 30% yield were dispersed in water at the desired concentration for future use, where the concentration was determined through the UV-Vis calibration curve of H_3_TATB solution.

#### b. HOF-BTB nanoparticles

30 mg H_3_BTB was dissolved in 2 mL DMF, and then 12 mL distilled water was added with stirring. After 5 min stirring, the products were collected through centrifuge 5 min at 12000 rpm (13523 x g), and washed with methanol and water 3 times, respectively. The yield was around 25%.

#### c. HOF-101 nanoparticles

Briefly, 30 mg H_4_TBAPy was dissolved in 3 mL DMF, and then dropwise added into 12 mL distilled water with stirring at 1000 rpm. After 5 min stirring, the nanoparticles were separated via centrifugation at 12000 rpm (13523 x g) for 5 min, and then washed with acetone, ethanol, and distilled water 3 times, respectively. The final products (yield: 95%) were resuspended with distilled water at the desired concentration for future use.

#### d. HOF-102 nanoparticles

10 mg H_4_PTTNA monomer was dissolved in 2 mL DMF, and then 8 mL methanol was added with stirring. After stirring for 5 min, the pallets were collected through centrifuge 5 min at 12000 rpm (13523 x g), and washed with methanol and water 3 times, respectively, with a yield 85%.

### Focused ultrasound (FUS) controlled scission of HOFs

Briefly, 2 mL fresh HOFs solution was loaded in glass vials, and placed on the surface of FUS transducer (Del Piezo Inc, DL-47, and Image Guided Therapy System). Ultrasound peak pressure was determined via hydrophone (Onda Corporation, HGL-0200) connected to a pre-amplifier (Onda Corporation, AG-2010) with a gain of 20 dB. The HOFs solution was irradiated under the FUS stimulation with different parameters. After that, 100 μL solution was extracted at a fixed time, and centrifuged at 8000 rpm for 5 min (Centrifuge 5430, Eppendorf), where the dissociated monomers or oligomers were separated and mainly located in the supernatant. The supernatant was extracted to do the UV-Vis tests (Eppendorf BioSpectrometer® Basic spectrophotometer) to determine the HOF dissociation percentage according to the following formula: HOFs dissociation percentage =Absorption of supernatantAbsorption of initial HOF solution 100%

### Theoretical calculation of the cohesive energy of HOFs and prediction model building

Density functional theory (DFT) calculations done with the Vienna Ab-Initio Package.^49,50^ Core electrons are described within the projected augmented wave framework; valence electrons are described with a plane wave basis set up to an energy cutoff of 400 eV.^51^ The generalized gradient approximation in the form of the Perdew, Burke and Ernzehof functional is used to model electronic exchange and correlation. Van der Waals interactions are calculated using the DFT-D2 method.^52^ Solvation energies are calculated using the VASPsol implicit solvation model.^53^

For each HOF, a monomer is isolated from the crystal structure and optimized to find its energy, *E_mono_*. The crystal structures are used to calculate the energy of the HOF, *E_HOF_*, except for BTB, where a fraction of the experimental crystal structure is used as an approximation. The dissociation of the HOF is modeled as reaction (1) where a constructional unit breaks from the bulk HOF state and becomes a dissociated dissolved monomer.

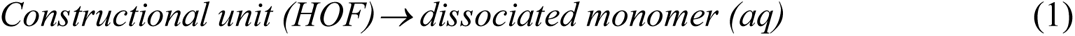

Given the experimentally measured dissociation percentage, *x,* at equilibrium, the dissociation equilibrium constant can be calculated using equation (2) below.

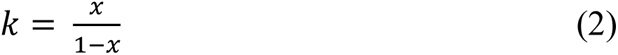

The cohesion energy *E_cohesion_* of each HOF in an aqueous environment is calculated using equation (3)

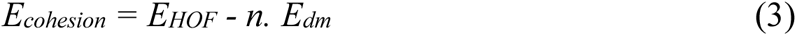

Where *E_HOF_* is the energy for a unit cell of the HOF, *n* is the number of constructional units in the unit cell, and *E_dm_* is energy of a dissociated monomer. For the crystal structures used in this study, *n* are 8, 2, 2, and 2 for HOF-TATB, HOF-BTB, HOF-101 and HOF-102, respectively. To characterize the hydrogen-bonding energy *E_HB_* and π-π interaction energy *E_π-π_* between two constructional units, a dimer bonded through hydrogen-bonding or π-π interaction is isolated from the crystal structure. The energy of the relaxed dimer is *E_dimer-HB_* and *E_dimer-π-π_* for the hydrogen-bonded dimer and the π-π bonded dimer, respectively. They are used to calculate *E_HB_* and *E_π-π_* using equations (4) and (5).

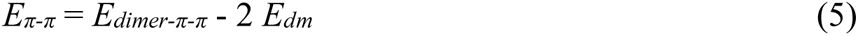

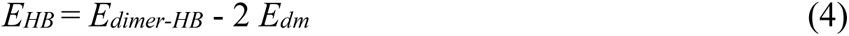

It’s worth noting that *E_HB_* is the energy per hydrogen-bonded dimer. One HOF-TATB building monomer can form 3 hydrogen bonds while one HOF-101 building monomer can form 4 hydrogen bonds. Therefore the total amount of hydrogen bonding energy a monomer can have is 1.5 *E_HB_* for HOF-TATB and 2 *E_HB_* for HOF-101. Structures of the dissociated monomers, hydrogen-bonded dimers, and π-π bonded dimers are shown in **Supplementary Table 4**.

### Preparation of drug-loaded HOF nanoparticles

2 mg of Rhodamine B (RB) or CNO was dissolved in 2 mL of a 5 mg/mL HOF-TATB or other HOFs nanoparticle solution. The mixtures were vibrated at 40 ℃ for 2 hours and then centrifuged at 12,000 rpm (13523 x g) for 5 minutes. The pellets were washed three times with distilled water to remove unloaded cargoes and then suspended in PBS. A 1 mL solution of nanoparticles was extracted for freeze-drying. Subsequently, the resulting powder (1 mg) was dissolved in 1 mL of water (pH 13) to completely release the cargo. The drug loading content was then measured using UV-Vis spectroscopy.

### In vitro ultrasound controlled drug uncaging

A solution of fresh drug-loaded HOF nanoparticles at a concentration of 1 mg/mL was loaded into glass vials and positioned on the ultrasound transducer. The solution was irradiated with FUS at a frequency of 1.5 MHz for a specific duration and under desired parameters. At predetermined time intervals, 100 μL of the solution was extracted and centrifuged at 8,000 rpm (6010 x g) for 5 minutes. The released drugs were found in the supernatant, and the percentage of drug release was calculated using UV-Vis spectroscopy.

### Long-term drug stability evaluation in HOF nanoparticles

Drug-loaded HOF nanoparticles at a concentration of 1 mg/mL were placed in glass vials and stored at room temperature. At specified time intervals, 100 μL of the solution was withdrawn and subjected to centrifugation at 8,000 rpm (6010 x g) for 5 minutes. The released drugs were detected in the supernatant, and the drug release percentage was determined using UV-Vis spectroscopy.

### In vitro calcium imaging

Calcium imaging was performed using primary cortical neurons that were transduced with AAV-9-hSyn-hM3D(Gq)-mCherry and pAAV-hSyn-GCaMP6s-WPRE-SV40. After 6 days of transduction, the neurons were fixed on a fluorescence microscope. The water balloon member of the FUS transducer was brought into contact with the neuron culture medium on top of the plate. Fresh TATB@CNO nanoparticles with a final concentration of 5 μg/mL CNO were added to the medium. Subsequently, FUS stimulation (1.08 MPa, 1.5 MHz, 10 s duration) was applied to trigger the release of CNO for neuron activation. Imaging videos were captured using a Leica DMi8 fluorescence microscope equipped with a 20X air objective, with a 5 ms exposure time, and green channel (Ex: 450-490 nm). The transient increase in green fluorescence (ΔF/F) was calculated by extracting the fluorescence time-series data of 100 neurons through manual segmentation and conversion of the video into grayscale using ImageJ software. The raw data was then processed using a custom MATLAB algorithm that detrends and normalizes the fluorescent time-series data through second-order polynomial curve fits and baseline maximum fluorescent value extraction, compensating for photobleaching effects.

### Stereotaxic injection of virus

All procedures designed according to the National Institute of Health Guide for the Care and Use of Laboratory Animals, approved by the Institutional Animal Care and Use Committee at the University of Texas at Austin (AUP-2021-00086, AUP-2021-00162), and were supported via the Animal Resources Center at the University of Texas at Austin. All the surgery tools were sterilized before the experiments. The hair was shaved before the injection, and the skin on the head was sterilized three times with 80% ethanol and iodophor. All the virus injections were conducted via the micro-injection system (World Precision Instruments, UMP3 Microinjection Syringe Pump) with 300 nL/min. After the injection, the needles remain inside the brain for at least 5 min to ensure the efficient diffusion of the virus, and then slowly withdraw in 5 min. The skin was closed with sutures after the injections. After the surgery, the animals were placed on the 37 ℃ heating pad and returned to the cage until fully recovered.

### Mouse model

C57BL/6 mice (20-26 g; 12-16 weeks old; Jackson laboratory) were used in our research. Mice were anesthetized with 2.5% isoflurane using anesthesia machine (Vaporizer Sales & Service Inc) and the head was fixed in stereotaxic frame (Kopf Stereotaxic Instruments). Each animal received a subcutaneous injection of meloxicam (5 mg/kg) and Ethiqa (3.25 mg/kg), and ophthalmic ointment was used to cover the eyes before surgery. For the photometry tests, 1200 nL AAV-9-hSyn-hM3D(Gq)-mCherry (100 µL at titer ≥ 1×10¹³ vg/mL)) and pAAV-hSyn-GCaMP6s-WPRE-SV40 (100 µL at titer ≥ 1×10¹³ vg/mL) mixture were unilaterally injected into the VTA, with the coordinates relative to bregma: anteroposterior (AP) −3.08 mm, mediolateral (ML) + 0.40 mm, and dorsoventral (DV) −5.0 mm.^34,48^ Of note, for the behavior tests, 1000 nL AAV-9-hSyn-hM3D(Gq)-mCherry solutions were unilaterally injected into the VTA. After 4 weeks, these mice were used for experiments.

### Rat model

3-4 months old Long-Evans Rats (Charles River) were used in our experiments. Rats were anesthetized with 5% isoflurane, and received a subcutaneous injection of meloxicam (2 mg/kg) and Ethiqa (0.65 mg/kg) before surgery, respectively. For the photometry tests, 1200 nL AAV-9-hSyn-hM3D(Gq)-mCherry and pAAV-hSyn-GCaMP6s-WPRE-SV40 mixture were unilaterally injected into the VTA, with the coordinates relative to bregma: anteroposterior (AP) −5.0 mm, mediolateral (ML) + 1.0 mm, and dorsoventral (DV) −8.6 mm.^54^ For the behavior tests, 1000 nL AAV-9-hSyn-hM3D(Gq)-mCherry was unilaterally injected into the VTA. After 4 weeks, these rats could be used for experiments.

### Mouse Photometry tests

The hM3D(Gq) and GCaMP6s transduced mice were used for the photometry tests. 2 μL of TATB@CNO (2 mg/mL) nanoparticles were unilaterally injected into the VTA with the same coordinates to the viral injection. Then, the optical fibers were implanted at the virus transduced VTA area (−3.08 mm AP, + 0.5 mm ML, −5.0 mm DV). After recovery for 2 days, the mouse head was head-fixed and the FUS (1.5 MHz, 1.40 MPa, 10 s duration) was used to irradiate the VTA area. FUS focus length was set to be 5 mm through control of the FUS water bubble. The signal was recorded through the R810 Dual Color Multi-Channel Fiber Photometry System (RWD life science).^55^

### Mouse FST evaluation

hM3D (Gq) transduced mice (20-24 weeks old) were used. A cylindrical tank (20 cm diameter x 30 cm height) was used for mice FST. The water level was around 15 cm, and the water temperature was around 23-25 ℃.^56^ The cylindrical tank was surrounded by a white background when we were ready to do the tests. On day 0, 2 μL of TATB@CNO (2 mg/mL) nanoparticles were unilaterally injected into the VTA (−3.08 mm AP, + 0.5 mm ML, −5.0 mm DV). After 2 days of recovery, the mouse was subjected to a 5 min pre-test without FUS stimulation on day 2. On day 3, FUS (1.5 MHz, 1.40 MPa, 20 s duration) was used to irradiate the VTA area through control of the FUS focus length (−3.08 mm AP, + 0.5 mm ML, −5.0 mm DV). After that, the mouse was placed in the FST container, and 5 min FST videos were recorded to analyze the immobility. Trajectory data was obtained from pre-built software integrated into the recorded FST videos using (R820 Tricolor Multi-Channel Fiber Photometry System, RWD). A custom-made algorithm in MATLAB was developed for FST data analysis. The difference in position between subsequent frames of the video was used to determine the displacement of the mice, which was then divided by the FPS of the video to determine the velocity. A threshold value of the velocity of 0.1 pixels/second was determined to be the optimal value for determining the state of mobility and immobility for mice in FST across groups. For every 5 second window, the time accrued between frames where the velocity is less than the threshold is accumulated and was used as proportion to the 5 second window to determine the immobility percentage over time. The total immobility time was determined similarly by accumulating the total time where the velocity between frames were below the threshold.

### Mouse Conditioned place preference tests

After the injection AAV (AAV9-hSyn-hM3D (Gq)-mCherry) for 4 weeks. 2 μL of TATB@CNO (2 mg/mL) nanoparticles were unilaterally injected into the VTA (−3.08 mm AP, + 0.5 mm ML, −5.0 mm DV) at day 0. Briefly, the two side boxes (30 cm X 60 cm X 30 cm) were made of plexiglas, and conjugated with one medium chamber (15 cm X 30 cm X 30 cm). One side box was covered with white and red striped papers, and another was covered with yellow and blue striped papers.^57^ On day 2, the mice were placed in the medium chamber to allow them freely to explore the apparatus for 15 min (pre-test). On day 3, mice were conditioned with 2 sessions. At first, the mice were irradiated with FUS (1.5 MHz, 1.40 MPa, 20 s duration), and then restricted in designated conditioned chambers for 30 min. In the second session around 3 h later, mice received the same treatment and were then restricted in designated conditioned chambers for 30 min. On day 4, the mice were placed and restricted in the medium chamber for 5 min. After that, the mice were allowed to freely move. The videos were recorded through a trail tracking camera. The data was analyzed through R810 Dual Color Multichannel Fiber Photometry with a behavior tracking system (RWD life science). For place preference tests, mice was chosen for tests only if the baseline preference for either side chamber is between 10%-70%, or for the medium chamber is <40%.^57^

### Rat Photometry tests

hM3D(Gq) and GCaMP6s transduced rats were used. 2 μL of TATB@CNO (10 mg/mL) nanoparticles were unilaterally injected into the VTA with similar coordinates (−5.0 mm AP, + 1.0 mm ML, −8.6 mm DV). Then, the optical fibers were implanted into a similar area. After recovery for 2 days, the rat head was fixed, and FUS (1.5 MHz, 2.45 MPa, 20s duration) was used to irradiate the VTA area. The FUS focus length was set to 9 mm through control of the FUS water bubble. The signal was recorded through the R810 Dual Color Multichannel Fiber Photometry System (RWD life science).

### Rat Conditioned place preference tests

After 4 weeks of expression of hM3D(Gq) in VTA neurons in rats, 2 μL of TATB@CNO (10 mg/mL) nanoparticles were unilaterally injected into the VTA (−5.0 mm AP, + 1.0 mm ML, −8.6 mm DV) at day 0. Similar to mice conditioned place preference tests, the two side boxes (30 cm X 60 cm X 30 cm) were made of plexiglass, but conjugated with one medium chamber (25 cm X 30 cm X 30 cm). On day 2, the rats were placed in the medium chamber, and stable for 5 min. After that, we opened the door to allow them freely to explore the apparatus for 15 min (pre-test). On day 3, rats were conditioned with 2 sessions. At first, the rats were irradiated with FUS (1.5 MHz, 2.45 MPa, 20s duration), and then restricted in designated conditioned chambers for 30 min. In the second session around 3 h later, rats received the same treatment and were then restricted to designated conditioned chambers for 30 min. On day 4, repeat the 2 pairing sessions similar to day 3. After two days of training, we started the tests on day 5, the rats were placed and restricted in the medium chamber for 5 min. After that, the rats were allowed to freely move for 15 min. The data analysis methods were similar to the mice conditioned place preference tests.

### c-fos staining in mice/rats brain sections

Specifically, mice/rats expressing hM3D (Gq) in VTA neurons and wild-type mice/rats were first subjected to sono-chemogenetics treatment. After 60 minutes, the mice/rats were anesthetized with ketamine (16 mg/kg) administered intraperitoneally. Following induction of deep anesthesia, perfusion was performed using PBS, followed by 4% paraformaldehyde. The brains were then extracted and stored in 4% paraformaldehyde at 4℃ overnight, and sliced using a vibrating blade microtome (Leica VT1200). Brain slices with a depth of 60 μm were washed with 0.3% Triton-X PBS (TBS) solution and subsequently blocked with 5% bovine serum albumin TBS solution for 30 minutes at room temperature. In the case of mouse brain sections, following the blocking step, the samples were incubated with rabbit anti-c-Fos antibody (ab222699, Abcam, 1:500)/mouse anti-tyrosine hydroxylase antibody (MA1-24654, Fisher Scientific, 1:1000)/0.3% Triton-X in PBS. The samples were then incubated at 4°C overnight and washed three times with TBS solution. Next, a mixture of TBS and secondary antibodies, including goat anti-rabbit Alexa Fluor 405 (ab175652, Abcam, 1:500), goat anti-mouse Alexa Fluor 488 (ab150113, Abcam, 1:1000), and Hoechst 33342 (17535, ATT Bioquest, 1:5000), was added and the slices were incubated for 2 hours at room temperature in a dark room. The slices were then washed three times with TBS, mounted on slides using mounting media (9990402, Fisher Scientific), and covered with a coverslip. Confocal images were obtained using a Zeiss 710 laser scanning microscope. For rat brain sections, the procedure was identical except for the use of rabbit anti-c-Fos antibody (ab289723, Abcam, 1:500) instead of ab222699. Detailed information on the antibodies used in this study can be found in **Supplementary Table 8**.

## Acknowledgment

TEM image acquisition was performed with the help of Michelle Mikesh at the Center for Biomedical Research Support Microscopy and Imaging Facility at UT Austin (RRID# SCR_021756). Dr. Huiliang Wang acknowledges funding support from the NIH Maximizing Investigators’ Research Award (National Institute of General Medical Sciences 1R35GM147408), the University of Texas at Austin Startup Fund, Robert A. Welsh Foundation Grant (No. F-2084-20210327) and Craig H. Neilsen Foundation Pilot Research Grant. We acknowledge BioRender.com for the figures drawing.

## Author Contributions

W.W., and H.W. designed the project. W.W. led this project and performed all the materials characterization, cell tests, animal tests and their analysis. Y.S., B.C. and Y.X. designed the HOFs and synthesized the materials. W.C. and G. H. conducted molecular simulation computing and discussed the data. K. T., I.P., and X.L., helped W.W. build animal models and animal behavior tests. J.J., J.H., A.L., and B. A. helped with animal behavior data analysis and immunohistology tests. All the co-authors contributed to the writing of the manuscript.

## Additional Information

Supplementary Information is available for this paper.

## References

1. Mirvakili, S. M. & Langer, R. Wireless on-demand drug delivery. Nature Electronics 4, 464– 477 (2021).

2. Boulatov, R. The liberating force of ultrasound. Nature chemistry 13 112–114 (2021).

3. Rwei, A. Y. et al. Ultrasound-triggered local anaesthesia. Nat Biomed Eng 1, 644–653 (2017).

4. Sebesta, C. et al. Subsecond multichannel magnetic control of select neural circuits in freely moving flies. Nat. Mater.21,951–958 (2022).

5. Bhansali, D. et al. Nanotechnology for Pain Management: Current and Future Therapeutic Interventions. Nano Today 39, 101223 (2021).

6. Duan, X. et al. Smart pH-sensitive and temporal-controlled polymeric micelles for effective combination therapy of doxorubicin and disulfiram. ACS Nano 7, 5858–5869 (2013).

7. Deisseroth, K. Optogenetics: 10 years of microbial opsins in neuroscience. Nat. Neurosci. 18, 1213–1225 (2015).

8. Wang, J. B., Aryal, M., Zhong, Q., Vyas, D. B. & Airan, R. D. Noninvasive Ultrasonic Drug Uncaging Maps Whole-Brain Functional Networks. Neuron 100, 728–738 (2018).

9. Huo, S. et al. Mechanochemical bond scission for the activation of drugs. Nat. Chem. 13, 131–139 (2021).

10. Bar-Zion, A. et al. Acoustically triggered mechanotherapy using genetically encoded gas vesicles. Nat. Nanotechnol. 16, 1403–1412 (2021).

11. Wang, C. et al. Ultrasound-responsive low-dose doxorubicin liposomes trigger mitochondrial DNA release and activate cGAS-STING-mediated antitumour immunity. Nat. Commun. 14, 3877 (2023).

12. Yao, Y. et al. Remote control of mechanochemical reactions under physiological conditions using biocompatible focused ultrasound. Proc. Natl. Acad. Sci. U. S. A. 120, e2309822120 (2023).

13. Airan, R. D. et al. Noninvasive Targeted Transcranial Neuromodulation via Focused Ultrasound Gated Drug Release from Nanoemulsions. Nano Lett. 17, 652–659 (2017).

14. Chen, H. & Hwang, J. H. Ultrasound-targeted microbubble destruction for chemotherapeutic drug delivery to solid tumors. J Ther Ultrasound 1, 10 (2013).

15. Kiessling, F. et al. Recent advances in molecular, multimodal and theranostic ultrasound imaging. Adv. Drug Deliv. Rev. 72, 15–27 (2014).

16. Rapoport, N. Drug-Loaded Perfluorocarbon Nanodroplets for Ultrasound-Mediated Drug Delivery. Adv. Exp. Med. Biol. 880, 221–241 (2016).

17. Shi, Z., Wu, J., Song, Q., Göstl, R. & Herrmann, A. Toward Drug Release Using Polymer Mechanochemical Disulfide Scission. J. Am. Chem. Soc. 142, 14725–14732 (2020).

18. Cravotto, G., Gaudino, E. C. & Cintas, P. On the mechanochemical activation by ultrasound. Chem. Soc. Rev. 42, 7521–7534 (2013).

19. Huo, S. et al. Mechano-nanoswitches for ultrasound-controlled drug activation. Adv. Sci. 9, e2104696 (2022).

20. Ghanem, M. A. et al. The role of polymer mechanochemistry in responsive materials and additive manufacturing. Nat. Rev. Mater. 6, 84–98 (2021).

21. Akbulatov, S. et al. Experimentally realized mechanochemistry distinct from force-accelerated scission of loaded bonds. Science 357, 299–303 (2017).

22. Li, J., Nagamani, C. & Moore, J. S. Polymer mechanochemistry: from destructive to productive. Acc. Chem. Res. 48, 2181–2190 (2015).

23. Chen, Y., Mellot, G., van Luijk, D., Creton, C. & Sijbesma, R. P. Mechanochemical tools for polymer materials. Chem. Soc. Rev. 50, 4100–4140 (2021).

24. Wu, M.-X. & Yang, Y.-W. Metal-Organic Framework (MOF)-Based Drug/Cargo Delivery and Cancer Therapy. Adv. Mater. 29, (2017).

25. Bhunia, S., Deo, K. A. & Gaharwar, A. K. 2D Covalent Organic Frameworks for Biomedical Applications. Adv. Funct. Mater. 30, 2002046 (2020).

26. Lin, R.-B. et al. Multifunctional porous hydrogen-bonded organic framework materials. Chem. Soc. Rev. 48, 1362–1389 (2019).

27. Yang, Y. et al. Ethylene/ethane separation in a stable hydrogen-bonded organic framework through a gating mechanism. Nat. Chem. 13, 933–939 (2021).

28. Lin, R.-B. & Chen, B. Hydrogen-bonded organic frameworks: Chemistry and functions. Chem 8, 2114–2135 (2022).

29. Li, Y.-L. et al. Record Complexity in the Polycatenation of Three Porous Hydrogen-Bonded Organic Frameworks with Stepwise Adsorption Behaviors. J. Am. Chem. Soc. 142, 7218– 7224 (2020).

30. Boesmans, W., Hao, M. M. & Vanden Berghe, P. Optogenetic and chemogenetic techniques for neurogastroenterology. Nat. Rev. Gastroenterol. Hepatol. 15, 21–38 (2018).

31. Kim, C. K., Adhikari, A. & Deisseroth, K. Integration of optogenetics with complementary methodologies in systems neuroscience. Nat. Rev. Neurosci. 18, 222–235 (2017).

32. Gomez, J. L. et al. Chemogenetics revealed: DREADD occupancy and activation via converted clozapine. Science 357, 503–507 (2017).

33. Szablowski, J. O., Lee-Gosselin, A., Lue, B., Malounda, D. & Shapiro, M. G. Acoustically targeted chemogenetics for the non-invasive control of neural circuits. Nat Biomed Eng 2, 475–484 (2018).

34. Rao, S. et al. Remotely controlled chemomagnetic modulation of targeted neural circuits. Nat. Nanotechnol. 14, 967–973 (2019).

35. Nagai, Y. et al. Deschloroclozapine, a potent and selective chemogenetic actuator enables rapid neuronal and behavioral modulations in mice and monkeys. Nat. Neurosci. 23, 1157– 1167 (2020).

36. Wang, B. et al. A novel mesoporous hydrogen-bonded organic framework with high porosity and stability. Chem. Commun. 56, 66–69 (2019).

37. Yin, Q. et al. An ultra-robust and crystalline redeemable hydrogen-bonded organic framework for synergistic chemo-photodynamic therapy. Angew. Chem. Weinheim Bergstr. Ger. 130, 7817–7822 (2018).

38. Zentner, C. A. et al. High surface area and Z’ in a thermally stable 8-fold polycatenated hydrogen-bonded framework. Chem. Commun. 51, 11642–11645 (2015).

39. Adamantidis, A. et al. Optogenetics: 10 years after ChR2 in neurons—views from the community. Nat. Neurosci. 18, 1202–1212 (2015).

40. Tye, K. M. et al. Dopamine neurons modulate neural encoding and expression of depression-related behaviour. Nature 493, 537–541 (2013).

41. Meng, Y., Hynynen, K. & Lipsman, N. Applications of focused ultrasound in the brain: from thermoablation to drug delivery. Nat. Rev. Neurol. 17, 7–22 (2021).

42. Wang, W. et al. Ultrasound-Triggered In Situ Photon Emission for Noninvasive Optogenetics. J. Am. Chem. Soc. 145, 1097–1107 (2023).

43. Lee, J.-U. et al. Non-contact long-range magnetic stimulation of mechanosensitive ion channels in freely moving animals. Nat. Mater. 20, 1029–1036 (2021).

44. Duque, M. et al. Sonogenetic control of mammalian cells using exogenous Transient Receptor Potential A1 channels. Nat. Commun. 13, 600 (2022).

45. Xian, Q. et al. Modulation of deep neural circuits with sonogenetics. Proc. Natl. Acad. Sci. U. S. A. 120, e2220575120 (2023).

46. Matsubara, T. et al. Author Correction: Remote control of neural function by X-ray-induced scintillation. Nat. Commun. 13, 1950 (2022).

47. Chen, R. et al. Deep brain optogenetics without intracranial surgery. Nat. Biotechnol. 39, 161–164 (2021).

48. Wu, X. et al. Tether-free photothermal deep-brain stimulation in freely behaving mice via wide-field illumination in the near-infrared-II window. Nat Biomed Eng 6, 754–770 (2022).

49. Kresse, G. & Hafner, J. Ab initio molecular dynamics for liquid metals. Phys. Rev. B Condens. Matter 47, 558–561 (1993).

50. Kresse, G. & Joubert, D. From ultrasoft pseudopotentials to the projector augmented-wave method. Phys. Rev. B Condens. Matter 59, 1758–1775 (1999).

51. Blöchl, P. E. Projector augmented-wave method. Phys. Rev. B Condens. Matter 50, 17953– 17979 (1994).

52. Grimme, S. Semiempirical GGA-type density functional constructed with a long-range dispersion correction. J. Comput. Chem. 27, 1787–1799 (2006).

53. Sundararaman, R. & Schwarz, K. Evaluating continuum solvation models for the electrode-electrolyte interface: Challenges and strategies for improvement. J. Chem. Phys. 146, 084111 (2017).

54. Witten, I. B. et al. Recombinase-driver rat lines: tools, techniques, and optogenetic application to dopamine-mediated reinforcement. Neuron 72, 721–733 (2011).

55. Zan, G.-Y. et al. Amygdalar κ-opioid receptor-dependent upregulating glutamate transporter 1 mediates depressive-like behaviors of opioid abstinence. Cell Rep. 37, 109913 (2021).

56. Can, A. et al. The mouse forced swim test. J. Vis. Exp. e3638 (2012).

57. Airan, R. D., Thompson, K. R., Fenno, L. E., Bernstein, H. & Deisseroth, K. Temporally precise in vivo control of intracellular signalling. Nature 458, 1025–1029 (2009).

