## Supplementary Information for "Ultrasound programmable hydrogen-bonded organic frameworks for sono-chemogenetics"

***Experimental Section/Methods***

***Materials and Instrumentation***

***Chemicals.*** Unless otherwise mentioned, all reagents and solvents were purchased from commercial sources and used as received without further purification. Deatils: (4-(methoxycarbonyl)phenyl)boronic acid, 1,3,6,8-tetrabromopyrene, palladium tetrakis(triphenylphosphine), potassium carbonate, 1,4-dioxane, sodium hydroxide （NaOH), potassium hydroxide (KOH), tetrahydrofuran (THF), concentrated hydrochloric acid (HCl), acetone, dimethylformamide (DMF), methanol, ethanol, palladium (II) acetate, methyl 6-bromo-2-naphthoate, sodium acetate (NaOAc), Palladium(II) chloride ($PdCl_2$), dichloromethane ($CH_2Cl_2$), magnesium sulfate ($MgSO_4$), 4,4,4',4',5,5,5',5'-octamethyl-2,2'-bi(1,3,2-dioxaborolane), dimethyl sulfoxide (DMSO), silica gel, potassium phosphate ($K_3PO_4$), hexane, ethyl acetate, aluminum chloride ($AlCl_3$), 1,3,5-triphenylbenzene, acetyl chloride, bromine, 1-propanol, 1-butanol, sodium bicarbonate, Sodium thiosulfate, 4-Methylbenzonitrile, tri-fluoro-methanesulfonic acid, aqueous ammonia, acetic acid, sulfuric acid (98%), chromium (VI)-oxide, acetic anhydride.

**Instruments**. Gas adsorption/desorption isotherms were measured by the volumetric method using a Micromeritics ASAP2020 surface area and pore analyzer. The powder X-ray diffraction patterns were collected by a Panalytical X’Pert powder diffractometer equipped with a Cu-sealed tube ($\lambda$ = 1.54184 Å) at 40 kV and 40 mA over the 2$\theta$ range of 5−40°. Bruker 400 MHz Avance III HD Nano was used for collections of $^1$H NMR spectra ($^1$H NMR, 400 MHz).

**Bioreagents**. pAAV-hSyn-hM3D(Gq)-mCherry was a gift from Bryan Roth (Addgene viral prep # 50474-AAV9; http://n2t.net/addgene:50474;RRID:Addgene_50474). (AAV-hSyn-GCaMP6s-WPRE-SV40 was a gift from Douglas Kim & GENIE Project (Addgene viral prep # 100843-AAV9; http://n2t.net/addgene:100843; RRID:Addgene_100843)[1] Clozapine N-oxide (CNO) dihydrochloride (water soluble) was ordered from Hello Bio, Rhodamine B was ordered from Sigma-Aldrich,

**Synthesis of the HOF-BTB building units 1,3,5-Tris(4-carboxyphenyl) benzene ($H_3$BTB) ligand.** 1,3,5-Tribromobenzene (1.00 g, 3.1 mmol), 4-ethoxycarbonylphenylboronic acid (2.22 g,

11.4 mmol) and $K_3PO_4$ (5.06 g, 23.8 mmol) were mixed in 1,4-dioxane (80 mL) and the mixture was degassed under argon for 10 min. [Pd($PPh_3$)$_4$] (0.03 g, 0.031 mmol) was added to the reaction mixture with stirring and the mixture was heated to 85 $^oC$ for 4 days under argon. The solution was evaporated to dryness and the residue was extracted with chloroform. The final product H3L was obtained by hydrolyzing the crude product with 2M aqueous NaOH, followed by acidification with concentrated HCl (1.18 g, 85 %).$^1$H-NMR (400 MHz, DMSO-d6): δ = 13.03 (s, 3H), 8.08 (d, 9H), 8,02 (d, 6H).

**Synthesis of HOF-TATB building unit 4,4',4''-(1,3,5-Triazine-2,4,6-triyl)-tribenzoic acid ($H_3$TATB) ligand.** 1, 4-Methylbenzonitrile (10.0 g, 86.0 mmol) was slowly added to tri-fluoro-methanesulfonic acid (20.9 mL, 236 mmol). After stirring at room temperature for 17 h, the red brown reaction mixture was neutralized with aqueous ammonia solution (2 M) with ice. The white precipitate was filtered and washed with a large amount of deionized water and acetone. The crude product was then recrystallized from toluene and dried under vacuum conditions. After that, 2,4,6-tris(4-methylphenyl)-1,3,5-triazine (8.1 g, 23.0 mmol, yield: 80 %) in the form of white crystals was obtained. Then, 2,4,6-tris(4-methylphenyl)-1,3,5-triazine (7.92 g, 22.5 mmol) was poured into a 500 mL flask, acetic acid (217 mL, 3.79 mol) and sulfuric acid (12.5 mL, 234 mmol) were added, the mixture solution was stirred, then chromium (VI)-oxide (20.5 g, 205 mmol) and acetic anhydride (13.7 mL, 145 mmol) were added carefully in small amounts (1.5 g and 1 mL, respectively). The dark-brown slurry was stirred at room temperature for 72 h, then poured into cold water (900 mL), well mixed, and then separated by centrifugation. The precipitates were collected, washed with water, and dissolved in aqueous sodium hydroxide solution (2 M, 500 mL). Unreacted starting material was removed by filtration, then the solution was acidified with hydrochloric acid (2 M) and washed with water to yield a slightly yellow crude product. After recrystallization from N, N-dimethylformamide and $H_2O$, 7.37 g (16.7 mmol, 74 %) white powder product was obtained. $^1$H-NMR (400 MHz, DMSO-d6): δ = 13.24 (s, 3H), 8.63 (d, 6H), 8,06 (d, 6H).

**Synthesis of HOF-101 building unit 1,3,6,8-Tetrakis (benzoic acid) pyrene (H4TBAPy) ligand.** A mixture of (4-(methoxycarbonyl) phenyl) boronic acid (5 g, 32.9 mmol), 1,3,6,8-tetrabromopyrene (2.85 g, 5.5 mmol), palladium tetrakis (triphenylphosphine) (0.1 g 0.09 mmol), and potassium carbonate (6 g, 44 mmol) in dry dioxane (100 mL) was stirred under the protection of $N_2$ for 72 h at 85 °C. The reaction mixture was poured into a solution of ice water and concentrated hydrochloric acid (v/v = 3:1). The organic phase was extracted with chloroform, and the combined extraction was dried over magnesium sulfate. After filtration, the solvent was removed under vacuum to give 3.44 g (4.6mmol)1,3,6,8-tetrakis(4-(methoxycarbonyl) phenyl) pyrene (yield: 84%). Then, 1g (17.8 mmol) KOH was added to a suspension of 1 g (1.465 mmol) of 1,3,6,8-tetrakis(4-(methoxycarbonyl) phenyl) pyrene in 100 mL of THF/dioxane/$H_2O$ (v/v = 5:2:2), and the mixture was stirred under reflux at 85 °C for 12 hours. The solvent was removed under vacuum, and then 100 mL $H_2O$ was added to the residue. The mixture (yellow clear solution) was stirred at room temperature for 2 h. The pH value was adjusted to 2 using concentrated HCl. The resultant yellow solids were collected by filtration and washed with water several times. After drying under vacuum, 0.88 g (1.29 mmol) product was obtained with a yield of 97%. $^1$H-NMR (DMSO-d6): δ= 13.10 (s, 4H), 8.20 (s, 4H), 8.15 (d, 8H), 8.08 (s, 2H), 7.85 (d, 8H).

**Synthesis of HOF-102 building unit 1,3,6,8-Tetra(6-carboxynaphthalen-2-yl) pyrene (H4PTTNA) ligand.** Dioxane (250 mL) was placed in a 500 mL three-necked round-bottom flask and purged with argon for 1.5 h. With argon purging and the use of a mechanical stirrer, tetrabromopyrene (5.0 g, 9.7 mmol), methyl 6-(pinacolboryl)-2-naphthoate (13.3 g, 42.5 mmol), potassium phosphate tribasic (16.5 g, 77.7 mmol), and tetrakis (triphenylphosphine)palladium (0.55 g, 0.48 mmol) were added. The reaction was heated to 90 °C for 72 h. The reaction mixture was allowed to cool to room temperature, and then 250 mL of water was added. The yellow solid was filtered using a glass Büchner funnel (medium frit) and washed with water (2×500 mL), followed by 500 mL of acetone. The filter flask was then emptied, and the solid was collected by passing hot chloroform (6×500 mL) through the frit and collecting thefiltrate. Purification by flash column chromatography (silica gel) afforded the L1-OMe product as a light yellow solid (5.28 g, 58%). Then, L1-OMe (5g, 5.3 mmol) was added to a 1000 mL single-necked flask, and dioxane (250 mL) was then added while stirring. A solution of potassium hydroxide (3.0 g, 53 mmol; 250 mL of water) was added, and the reaction mixture was heated to reflux while being vigorously stirred for 18 h (at this point, a clear solution was observed). The reaction was allowed to cool to room temperature. The organic solvent was removed using rotary evaporation, and 500 mL of water was added to dissolve the solid obtained. Concentrated HCl was added to the solution dropwise with stirring until the solution reached pH 1. After being stirred for an additional hour, the yellow precipitate was collected via centrifugation and washed with water (3×50 mL). The final solid product was recrystallized from DMF, filtered, and dried (3.6 g, 68%).$^{1}$H-NMR (DMSO-d6): δ= 13.14 (s, 4H), 8.71 (s, 4H), 8.49 (s,4H), 8.32 (d, 4H), 8.29 (s, 6H), 8.11 (d, 4H), 8.08 (d, 4H), 7.94 (d, 4H).

**Thermolysis of HOFs.** 1 mg/mL fresh HOFs solution was loaded into a centrifuge tube, and heated with a heater (Fisherbrand™ Isotemp™ Digital Dry Baths/Block Heaters) for 5 min at the designed temperature. After that, the solution was centrifuged at 8000 rpm for 5 min, and the supernatant was extracted to do the UV-Vis tests to determine the HOF dissociation percentage.

**Cell viability tests**. Human embryonic kidney 293 (HEK293T) cells were seeded into 96 well plates coated with 10 μg/mL Poly-L-Ornithine solution and cultured in DMEM with 10% fetal bovine serum at 37 °C under 5% $CO_2$. Then, complete medium containing TATB nanoparticles at various concentrations were added into the plate when the confluency was around 90%. After incubation for 24 h, Cell-Titer blue reagent (10 μL, Promega Corporation) was added into each well. Microplate reader (BioTek Synergy H1, 560ex/590em nm) was used to measure the cell viability after 4 h incubation according to the following formula:

$$\text{Cell viability (\%)} = \frac{\text{Fluorescence intensity of sample}}{\text{Fluorescence intensity of control}} \times 100\%$$

**Hemolysis tests of TATB nanoparticles.** According to the previous method, 0.3-0.5 mL fresh blood was extracted from the mouse heart, and then washed 3 times with 5 mL PBS after centrifuge at 8000 rpm for 5 min. The red blood cells (RBCs) were suspended with 2 mL PBS. To evaluate the biocompatibility of nanoparticles, 1 mL TATB nanoparticles at different concentrations were mixed with 30 μL RBCs. The mixture was incubated at 37 ℃ for 2 h, and then centrifuged at 12000 rpm for 5 min. The supernatant was extracted out for UV-Vis tests at absorption 541 nm. Notably, 30 μL RBCs were added to 1 mL distilled water as positive control or 1 mL PBS as negative control. The hemolysis percentage was calculated through the following formula:

$$\text{Hemolysis percentage (\%)} = \frac{\text{UV} - \text{Vis absorbance of sample}}{\text{UV} - \text{Vis absorbance of positive control}} \times 100\%$$

**Primary neuron culture**. C57BL/6 mice (8 weeks old, weighing 20-26 g; Jackson Laboratory) were utilized for our research. Primary cortical neurons were prepared from mice at embryonic day 15.5. Initially, 24-well cell culture plates were coated with poly-l-ornithine (0.2 mg/mL) and incubated at 37 °C for 2 hours. The plates were then washed three times with PBS and pre-warmed in a cell incubator for 15 minutes prior to use. Dissociated neural cells were plated on the coated plates and cultured in neurobasal medium supplemented with B27, glutamine, penicillin, and streptomycin. The neurons were incubated at 37 °C under 7% $CO_2$. After 2 days of incubation, a glial inhibitor, 5-fluoro-2'-deoxyuridine (0.1 mM), was added to the culture medium. Following 4 days of incubation, 1 μL of AAV-9-hSyn-hM3D(Gq)-mCherry and 1 μL of pAAV-hSyn-GCaMP6s-WPRE-SV40 were transfected into the neurons. Calcium imaging experiments were conducted after an additional 6 days of incubation.

**Ultrasound power heatmap determination in mouse and rat brain**. 57BL/6 mice (20-26 g; 12-16 weeks old; Jackson Laboratory) and Long-Evans rats (200-400 g; 12-16 weeks old; Charles River) were used in this study. The animals were sacrificed, and their heads were collected. The heads were then perfused with a solution of 4% paraformaldehyde (PFA) over a period of 24 hours to preserve the tissue. After perfusion, the heads were sliced into different thicknesses. To assess the ultrasound transmission efficiency in the head, a transducer (Image Guided Therapy System) was positioned in front of the head sections, and ultrasound gel (Aquasonic 100 Ultrasound Gel) was applied to ensure good contact between the transducer and the head sections. A hydrophone (HGL-0200, Onda Corporation) connected with a data acquisition oscilloscope (2204A-D2, Pico Technology) was placed behind the sections to accurately record the signals. The voltage measurements obtained from the hydrophone were then converted to absolute pressure values using the calibration specifications provided by the hydrophone vendor.

**Iba1, GFAP and Caspase-3 Staining.** Following the *in vivo* sono-optogenetics procedures, mice were subjected to anesthesia via intraperitoneal injection of ketamine (16 mg/kg) after 14 days. Perfusion was then performed using PBS and 4% paraformaldehyde. The brain was extracted and stored in 4% paraformaldehyde overnight at 4 °C and sliced using a vibrating blade microtome. Brain slices with a depth of 60 μm were washed with TBS solution and blocked with 5% bovine serum albumin TBS solution for 30 minutes at room temperature. Subsequently, the blocking buffer was replaced by rabbit anti-Iba1 antibody (013-27691, Wako Chem, 1:500)/TBS or rabbit anti-Cleaved Caspase-3 antibody (9661, Cell Signaling Tec., 1:500)/TBS or rabbit anti-GFAP antibody (13-0300, Invitrogen, 1:500)/TBA After incubation overnight at 4 ℃, the slices were washed three times with TBS solution. Fresh TBS solution containing the secondary antibody Donkey anti-rabbit Alexa Fluor 594 (A32754, Invitrogen, 1:500) and Hoechst 33342 (17535, ATT Bioquest, 1:5000) was added and incubated for 2 hours at room temperature in a dark room. Finally, the slices were washed three times with TBS, mounted on slides with mounting media, and covered with a coverslip. Fluorescence images were obtained using a Leica DMi8 fluorescence microscope.

**H&E staining.** Following the *in vivo* sono-optogenetics procedures, the mice were anesthetized with i.p. injection of ketamine after 14 days, and perfused with PBS and 4% paraformaldehyde. The brains were extracted, fixed in 4% paraformaldehyde overnight at 4 ℃, and sliced into 10 μm sections using a vibrating blade microtome. H&E staining was performed on the brain sections according to previously described protocols. The stained sections were then mounted onto slides with mounting media and covered with a coverslip. Images were captured using a Nikon Eclipse Ni Compound Light Microscope. Detailed information on the antibodies used in this study can be found in **Table S8**.

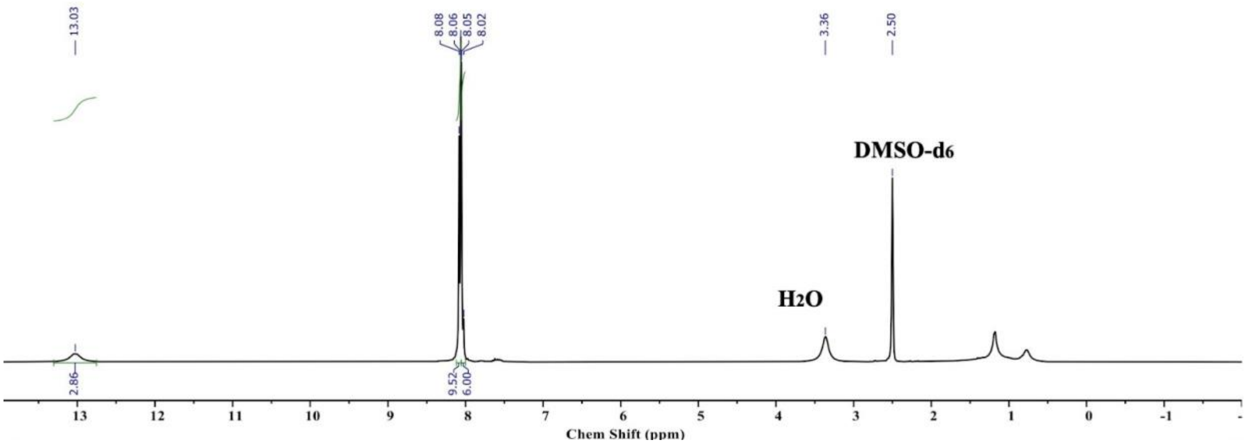


**Supplementary Fig. 1.** The [1]H NMR spectrum of HOF-TATB build unit ($H_3TATB$) ligand.

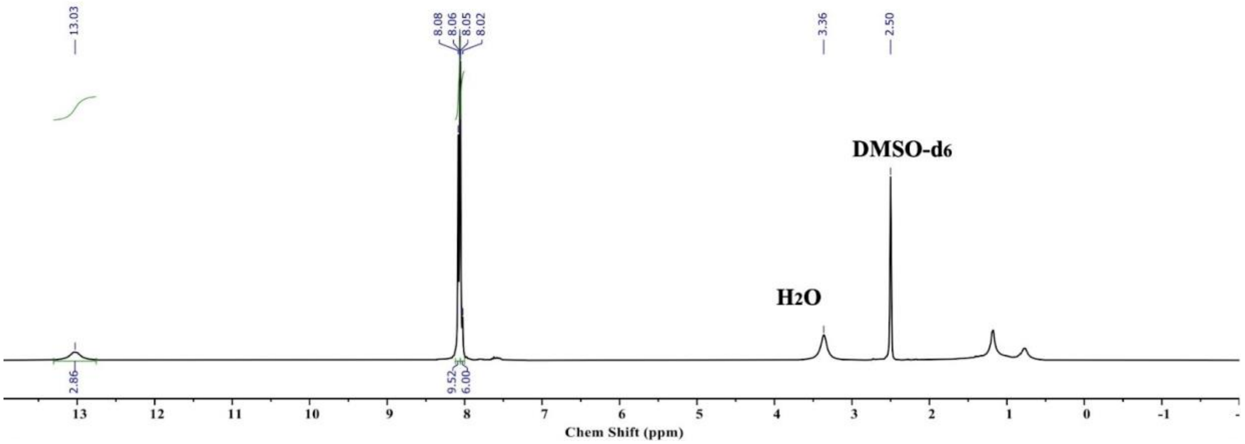


**Supplementary Fig. 2.** The [1]H NMR spectrum of HOF-BTB build unit ($H_3BTB$) ligand.

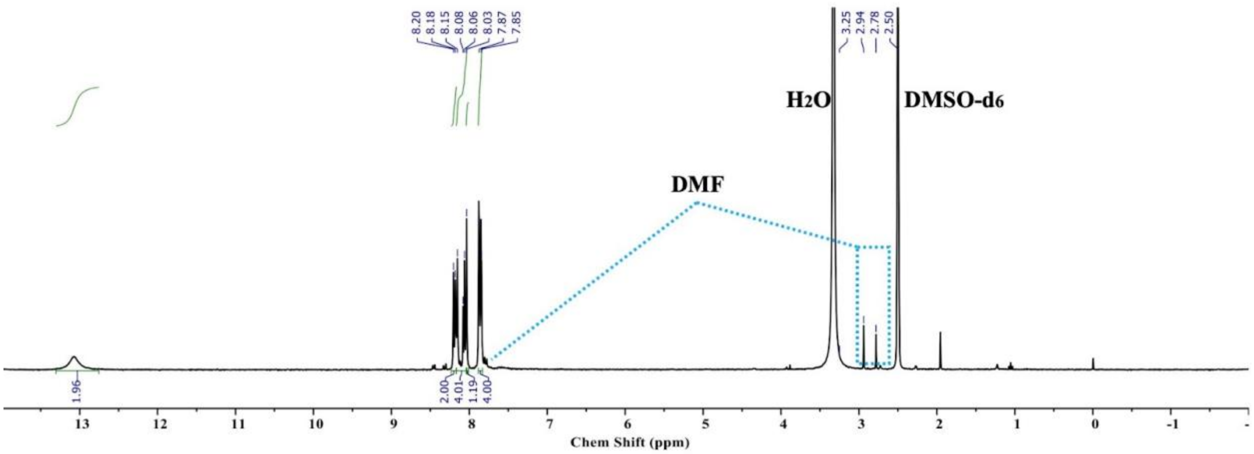


**Supplementary Fig. 3.** The [1]H NMR spectrum of HOF-101 build unit ($H_4$TBAPy) ligand.

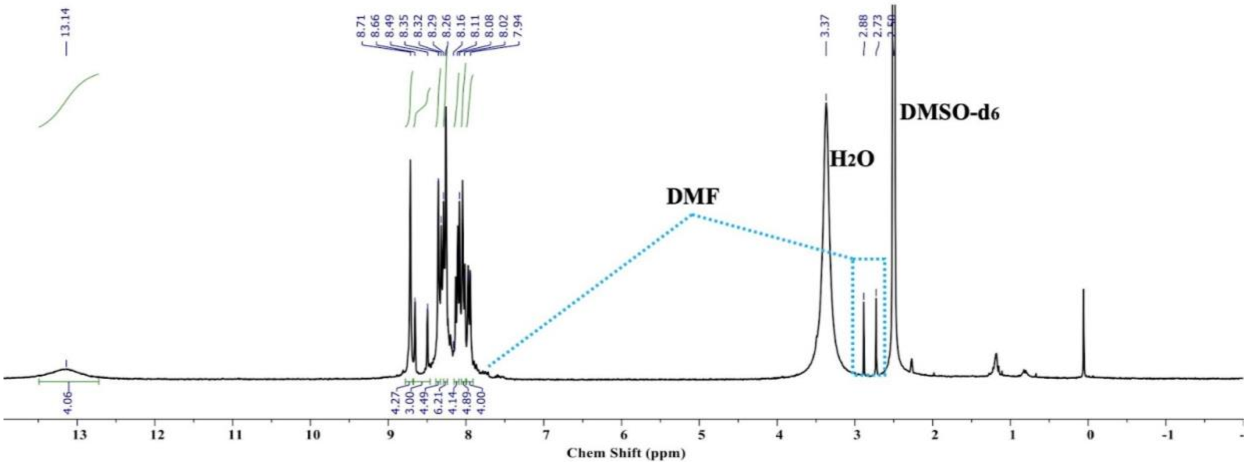


**Supplementary Fig. 4.** The [1]H NMR spectrum of HOF-102 build unit ($H_4$PTTNA) ligand.

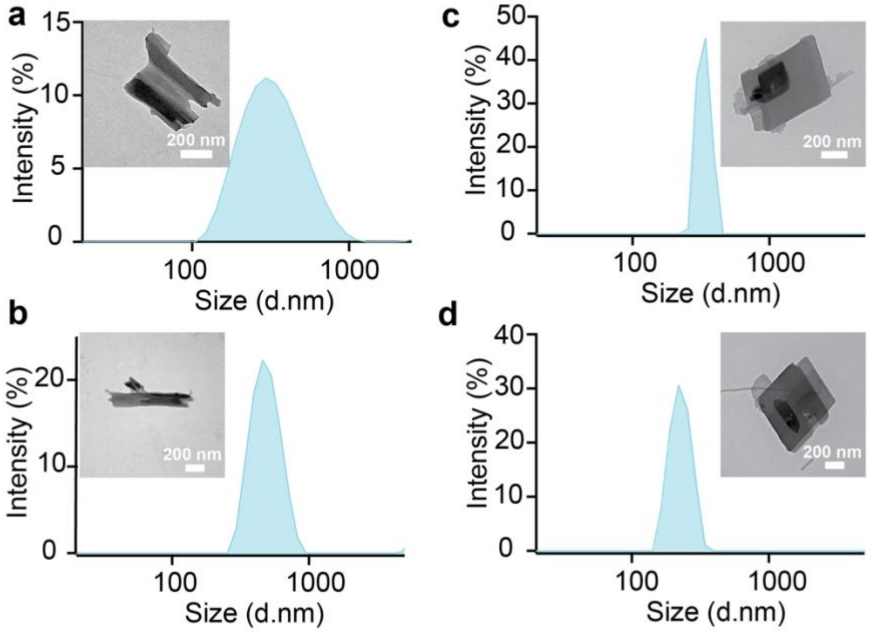


**Supplementary Fig. 5. TEM images and hydrodynamic size distribution measured by dynamic light scattering** of (a) HOF-TATB nanoparticles, (b) HOF-BTB nanoparticles, (c) HOF-101 nanoparticles, and (d) HOF-102 nanoparticles.

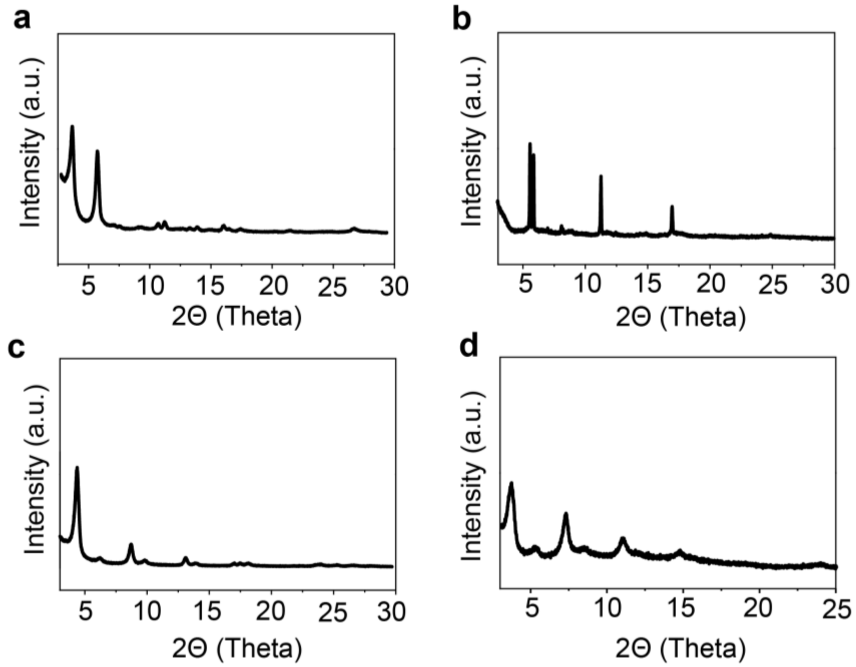


**Supplementary Fig. 6. The X-ray diffraction (XRD) tests of nanosized HOFs,** (a) HOF-TATB nanoparticles, (b) HOF-BTB nanoparticles, (c) HOF-101 nanoparticles, (d) HOF-102 nanoparticles.

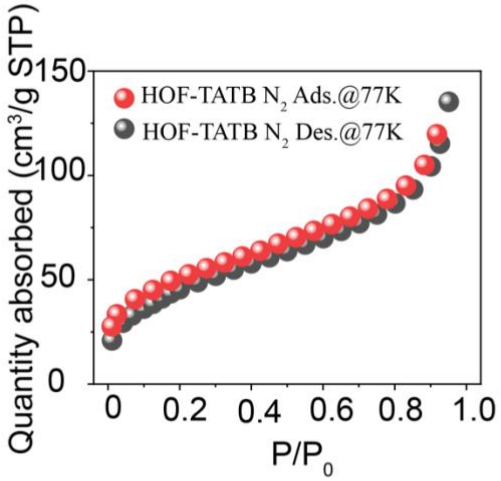


**Supplementary Fig. 7. The porosity characterization of HOF-TATB nanoparticles.** Single-component sorption isotherms of nitrogen at 77 K of nanosized HOF-TATB. BET Surface Area: 166.5 $m^2$ $g^{-1}$, Langmuir Surface Area: 292.11 $m^2$ $g^{-1}$.

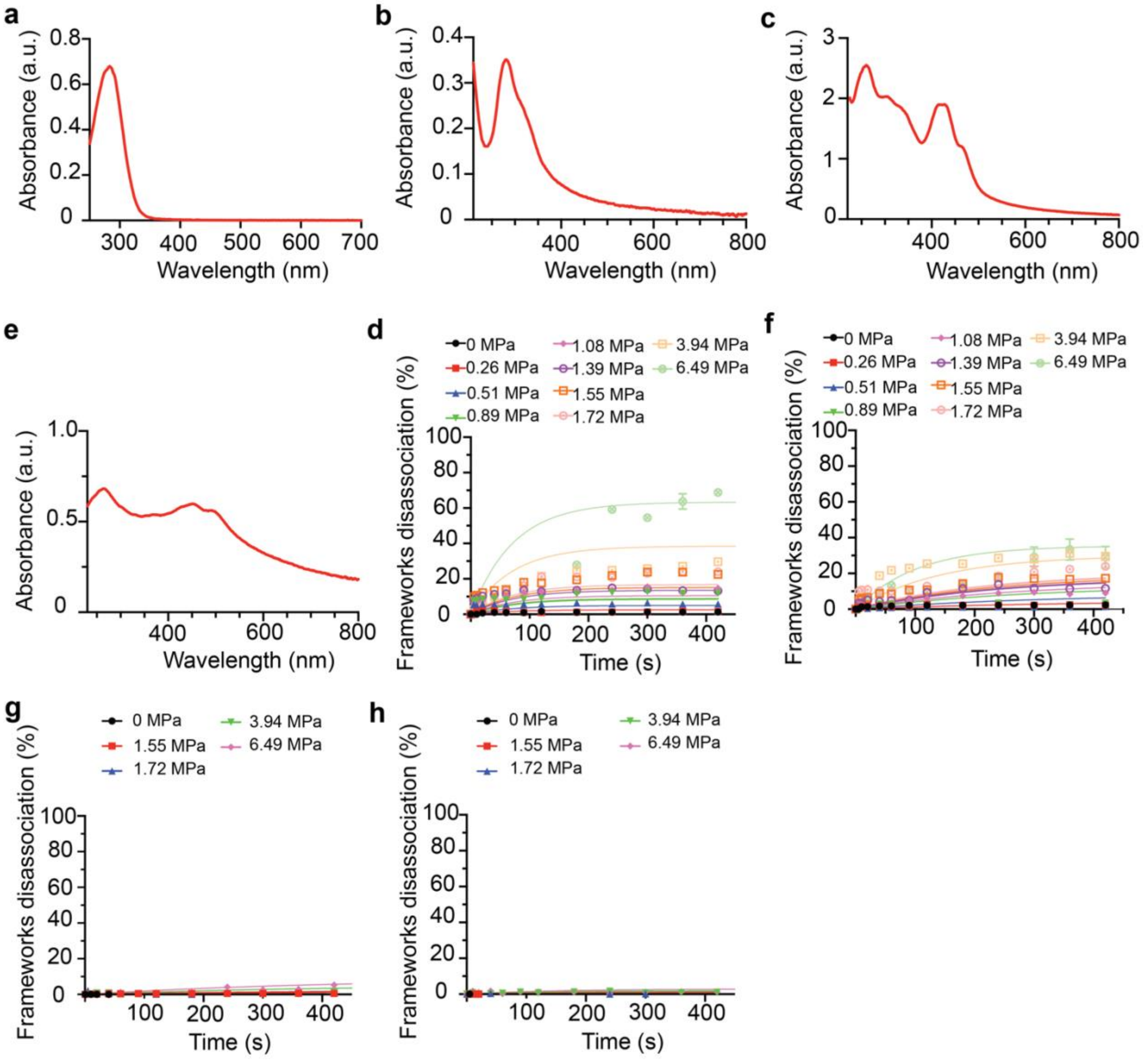


**Supplementary Fig. 8. FUS triggered dissociation of HOFs.** The HOFs nanoparticles were irradiated with different ultrasound powers, and the dissociation of HOFs was determined through UV-Vis tests. The characteristic spectrum of (a) HOF-TATB, (b) HOF-BTB, (c) HOF-101, (d) HOF-102. The dissociation curves of (e) HOF-TATB, (f) HOF-BTB, (g) HOF-101, (h) HOF-102 at different ultrasound peak pressures.

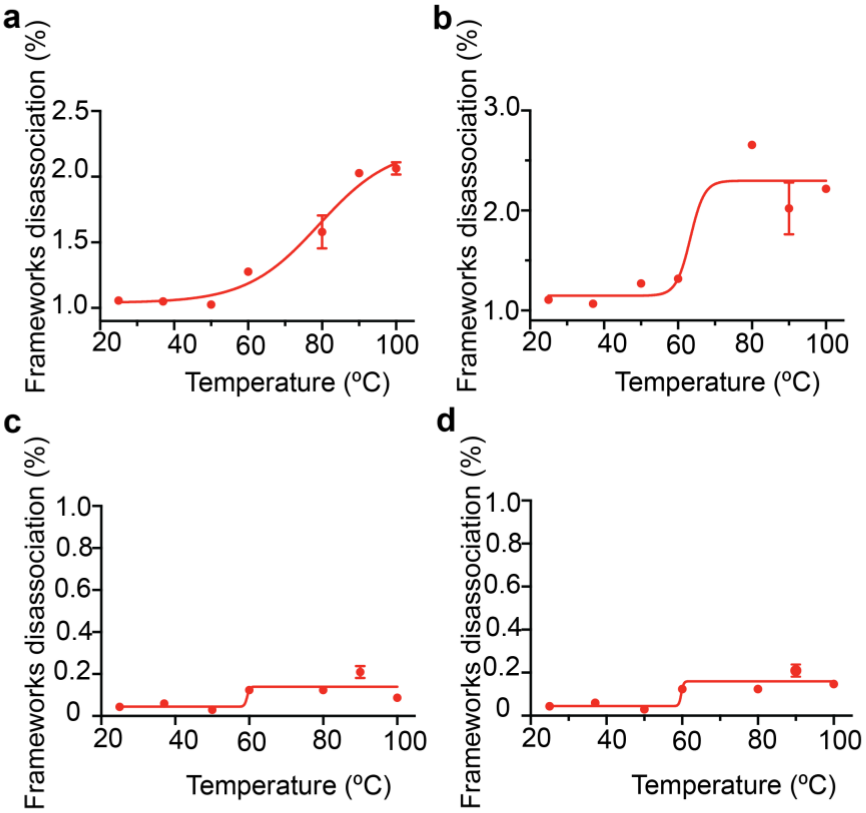


**Supplementary Fig. 9. The thermal dissociation tests of HOFs,** (a) HOF-TATB, (b) HOF-BTB, (c) HOF-101, (d) HOF-102. The HOFs nanoparticles were incubated at different temperatures for 5 min. After that, the HOFs solution was extracted out and centrifuged, the supernatant was conducted to do UV-Vis tests for HOFs dissociation determination. The thermal dissociation occurred around 60 °C. Only around a 2% increase was observed at HOF-TATB and HOF-BTB, and no thermal dissociation was observed in HOF-101 and HOF-102 the temperature at 100 °C.

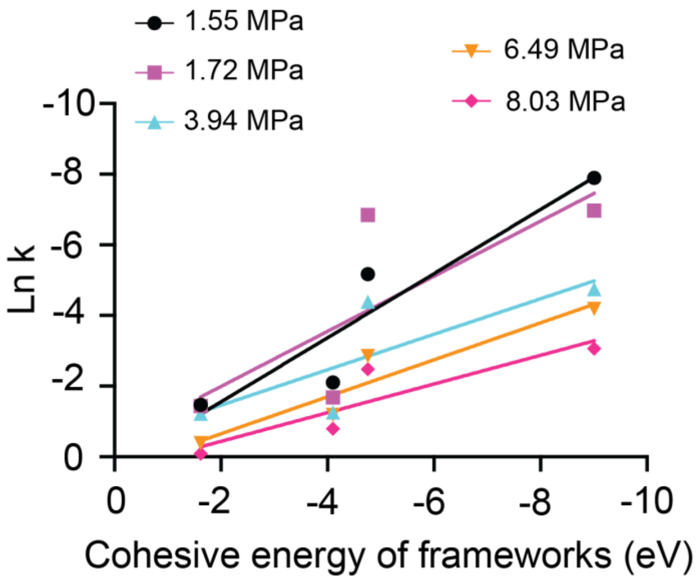


**Supplementary Fig. 10. A linear model fits the relationship between the $E_{cohesive}$ of HOFs and the Ln[k] at fixed $E_{us}$.** With 1.55, 1.72, 3.94, 6.49, 8.04 MPa peak pressure, Ln[k] of HOF-TATB, HOF-BTB, HOF-101, and HOF-102 correlates to their cohesion energy linearly, respectively.

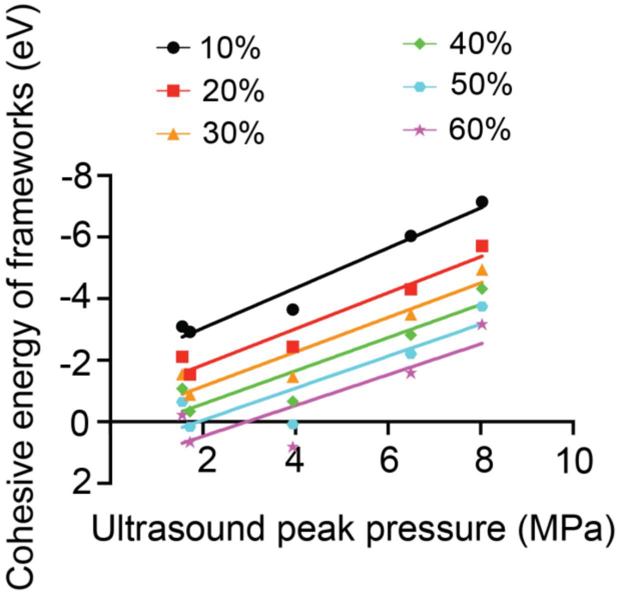


**Supplementary Fig. 11. When Ln[k] is held constant, a linear correlation is observed between the ultrasound peak pressure and the cohesive energy of HOFs.** To achieve a targeted 10%, 20%, 30%, 40%, 50% and 60% dissociation of HOFs at a fixed ultrasound peak pressure, it is possible to calculate the corresponding $E_{cohesive}$ of HOFs using the established linear relationship.

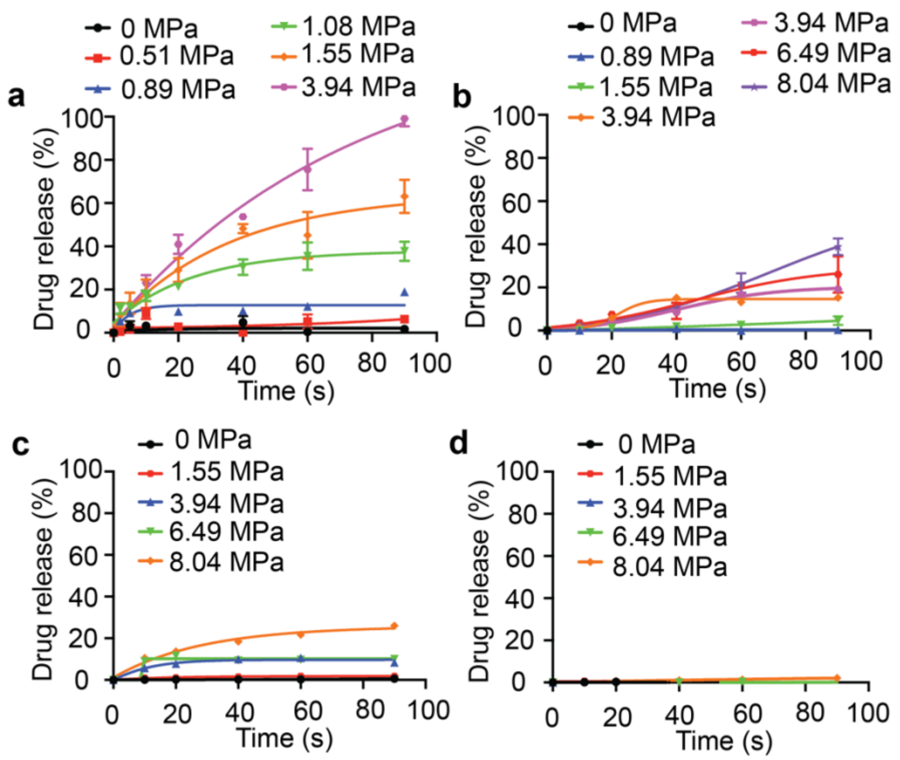


**Supplementary Fig. 12**. **Ultrasound triggered drug release (**mean ± SEM, n=3 independent samples**) from different HOF nanoparticles.** The fluorescence dye Rhodamine B (RB) was first loaded into the HOF nanoparticles. After that, the ultrasound irradiated the RB-loaded nanoparticles with different power densities. At fixed time points, the solution was taken out and centrifuged. The released RB concentration was determined via UV-Vis from the supernatant.

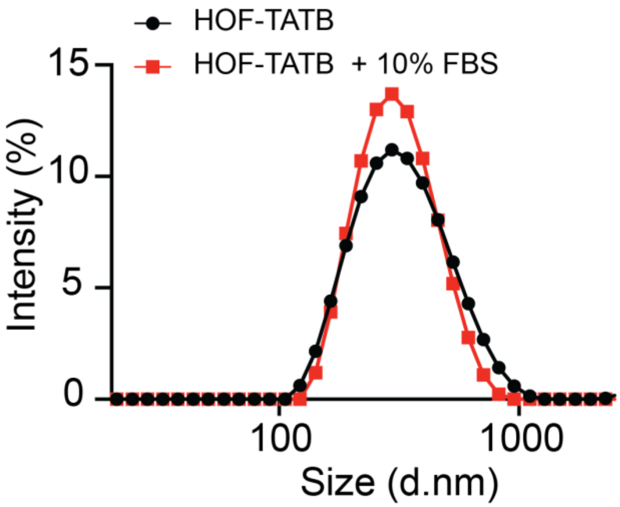


**Supplementary Fig. 13. The biostability evaluation of HOF-TATB nanoparticles.** The hydrodynamic size distribution was measured by dynamic light scattering with or without 10% FBS.

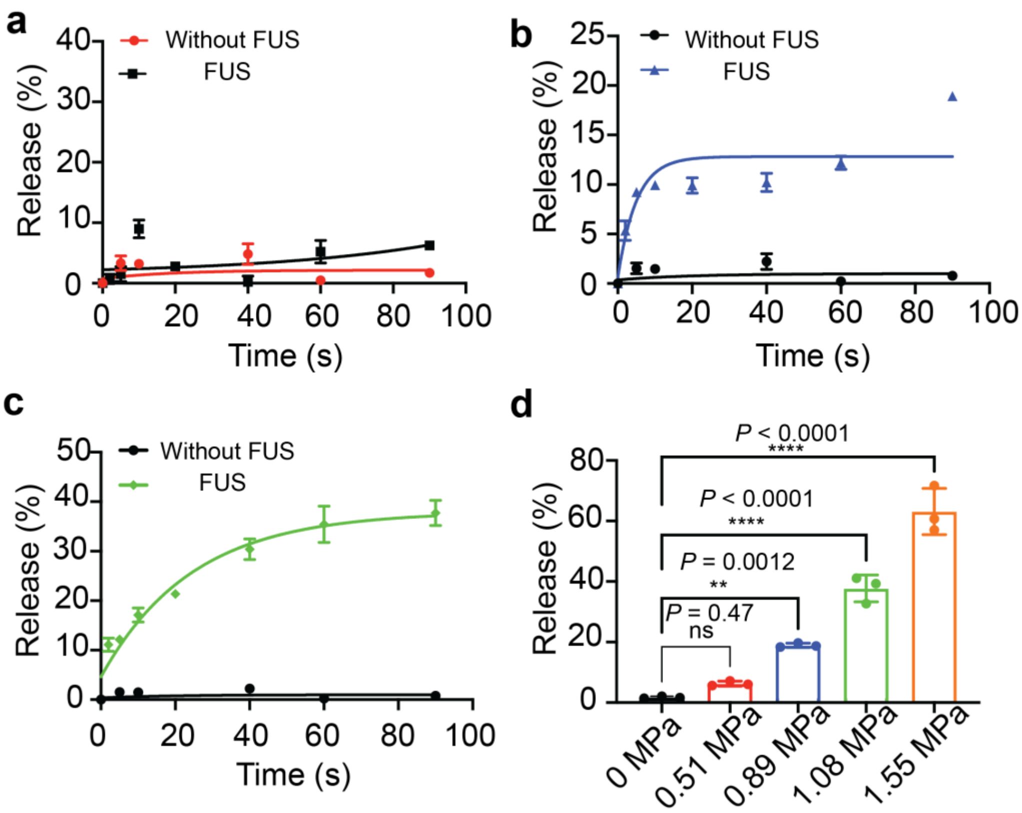


**Supplementary Fig. 14. Ultrasound triggered drug release (**mean ± SEM, n=3 independent samples**) from HOF-TATB.** The fluorescence dye Rhodamine B (RB) was loaded into the HOF-TATB nanoparticles (TATB@RB) at first. After that, the TATB@RB nanoparticles were irradiated by the ultrasound with different power densities, including (a) 0.51 MPa, (b) 0.89 MPa, and (c) 1.08 MPa. (d) The quantification of drug release percentage without ultrasound and with ultrasound for 90 s. Mean ± SEM, n ≥ 3 independent samples. One-way ANOVA and Dunnett's multiple comparison tests ($P \geq 0.05$ (ns), * $0.01 \leq P < 0.05$, ** $0.001 \leq P < 0.01$, **** $P < 0.0001$).

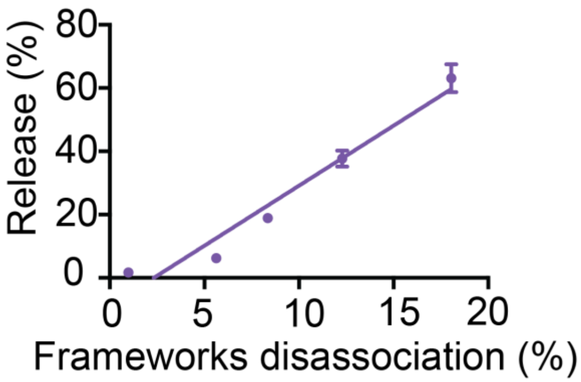


**Supplementary Fig. 15.** Ultrasound-triggered dye release at different frameworks disassociation percentage of the HOF-TATB nanoparticles. Mean ± SEM, n=3 independent samples.

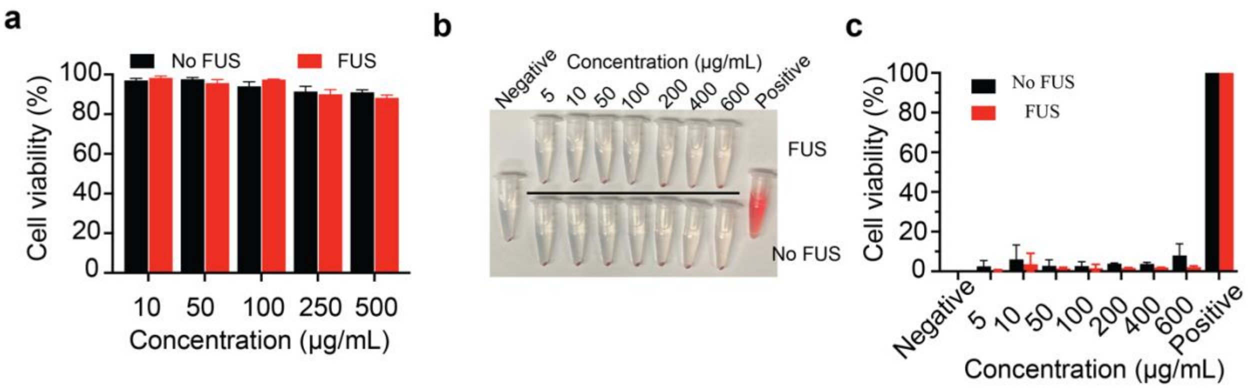


**Supplementary Fig. 16. The *in vitro* biosafety evaluation of HOF-TATB nanoparticles.** (a) The cell viability tests of HOF-TATB nanoparticles in human embryonic kidney 293 (HEK-293) cells. Mean ± SEM; at least 3 times independent tests ($n \geq 3$). The hemolysis tests of HOF-TATB nanoparticles, (b) photograph and (c) hemolysis statistical analysis, mean ± SEM; at least 3 times independent tests ($n \geq 3$).

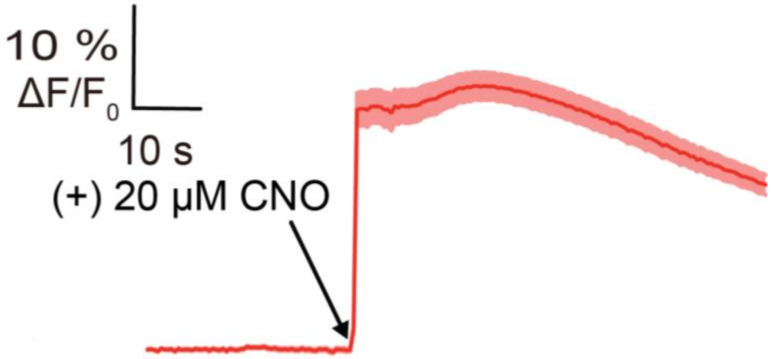


**Supplementary Fig. 17. Normalized GCaMP6s fluorescence intensity increase ($\Delta F/F_0$) from 100 of hM3D(Gq) and GCaMP6s expressing neurons in response to free CNO (20 μM).** Solid line, mean; shade area, SEM; at least 3 times independent tests ($n \geq 3$).

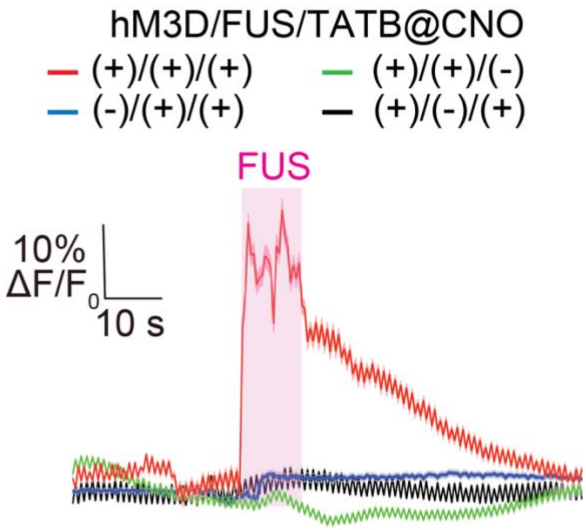


**Supplementary Fig. 18. Normalized GCaMP6s fluorescence intensity from 100 neurons in different experimental conditions.** The experiments were independently repeated at least three times. $F_0$ was the mean fluorescence intensity of baseline, hM3D(Gq)$^+$ means neurons with hM3D(Gq) expression, hM3D(Gq)$^-$ means neurons without hM3D(Gq) expression, FUS$^+$ means the neurons that are irradiated by the focused ultrasound and FUS$^-$ means the neurons without focused ultrasound irradiation, TATB@CNO$^+$ means HOF-TATB nanoparticles loaded with CNO is added to the neurons, TATB@CNO$^-$ means only HOF-TATB nanoparticles are added to treat the neurons. OFF, ultrasound is off. ON, ultrasound is on (1.5 MHz, 1.08 MPa, pulse 10 s).

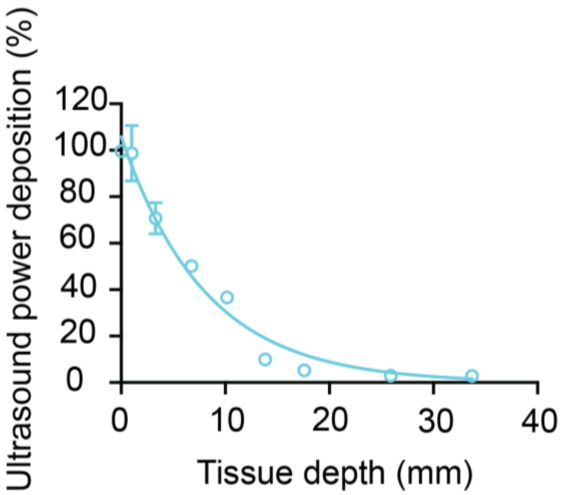


**Supplementary Fig. 19. Ultrasound power deposition in the tissue.** To determine the ultrasound power transfer efficiency in the tissue, pork skin with different depths was first placed on the FUS transducer (1.5 MHz, 2.40 MPa primary ultrasound power), and the ultrasound power was detected by the hydrophone. The results determined that the 1.5 MHz ultrasound could achieve around 20 mm tissue penetration, and the ultrasound power transfer efficiency was still at 37% even at 10 mm tissue depth.

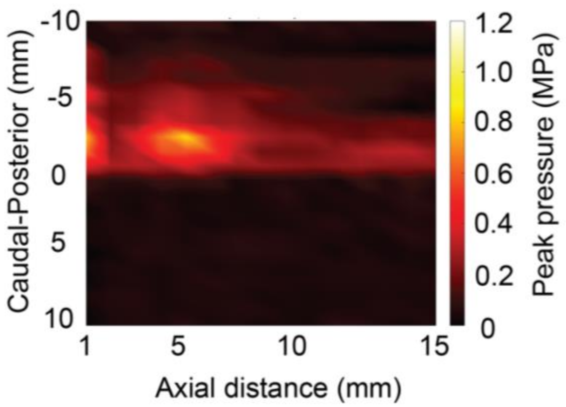


**Supplementary Fig. 20. The *in vivo* ultrasound power transfer in the mouse head with FUS focus length of 5 mm.** The ultrasound power heatmap in the mouse head shows that around 0.9 MPa was delivered to the mouse VTA when 1.40 MPa primary ultrasound power was used.

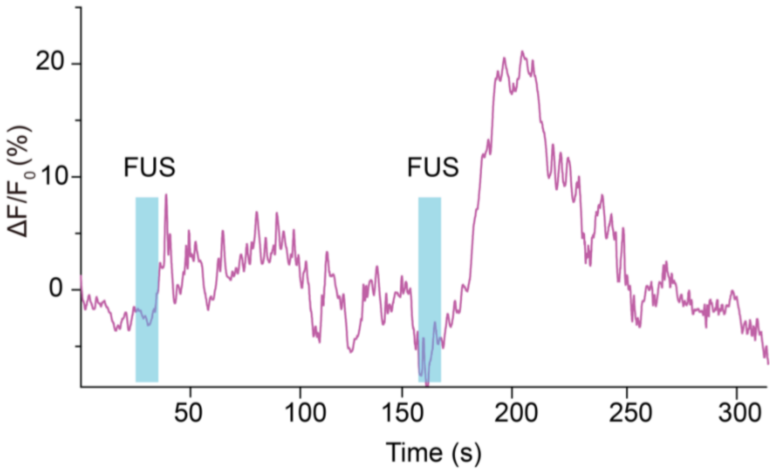


**Supplementary Fig. 21. Normalized GCaMP6s fluorescence intensity change (ΔF/$F_0$) in mice VTA under the repeated sono-chemogenetic stimulation after 5 days injection.** Multiple increases in GCaMP6s signal were recorded in mouse VTA under the repeated irradiation of sono-chemogenetic stimulation. FUS (1.5 MHz, 1.40 MPa, pulse 10 s), TATB@CNO nanoparticles (2 μg/mL). The blue area represents the FUS irradiation.

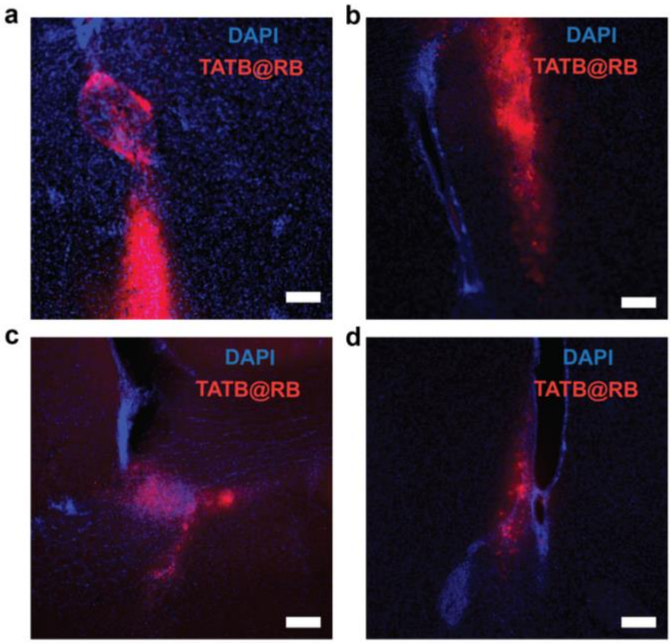


**Supplementary Fig. 22. *In vivo* stability evaluation of HOF-TATB nanoparticles.** Rhodamine B (RB) dye-loaded HOF-TATB nanoparticles (TATB@RB) were injected into the mouse VTA. 50 μm thick brain slices were prepared 1 day (a), 3 days (b), 2 weeks (c) and 3 weeks (d) following injection. **a-d**, fluorescence images of TATB@RB injection sites. Scale bar: 50 μm. Blue signal is DAP and, red signal is TATB@RB.

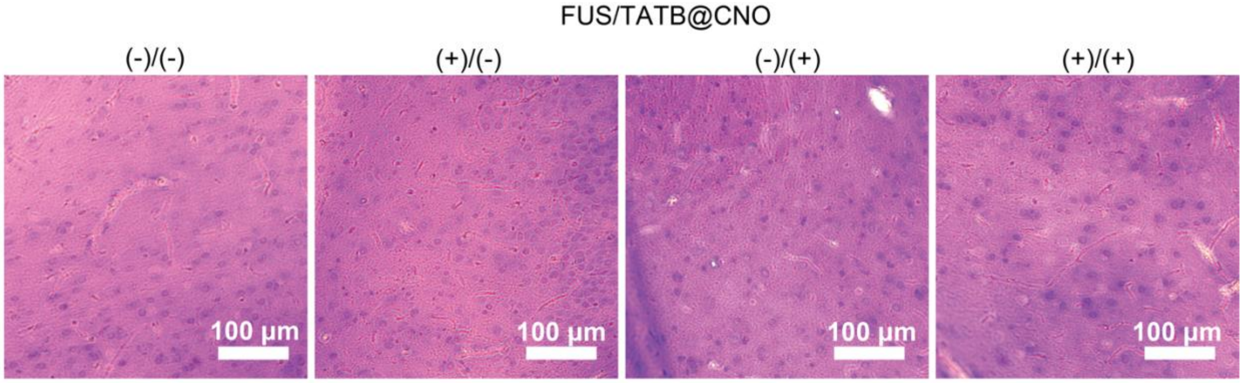


**Supplementary Fig. 23.** *In vivo* biosafety evaluation by H&E staining after sono-chemogenetics, scale bar: 100 μm.

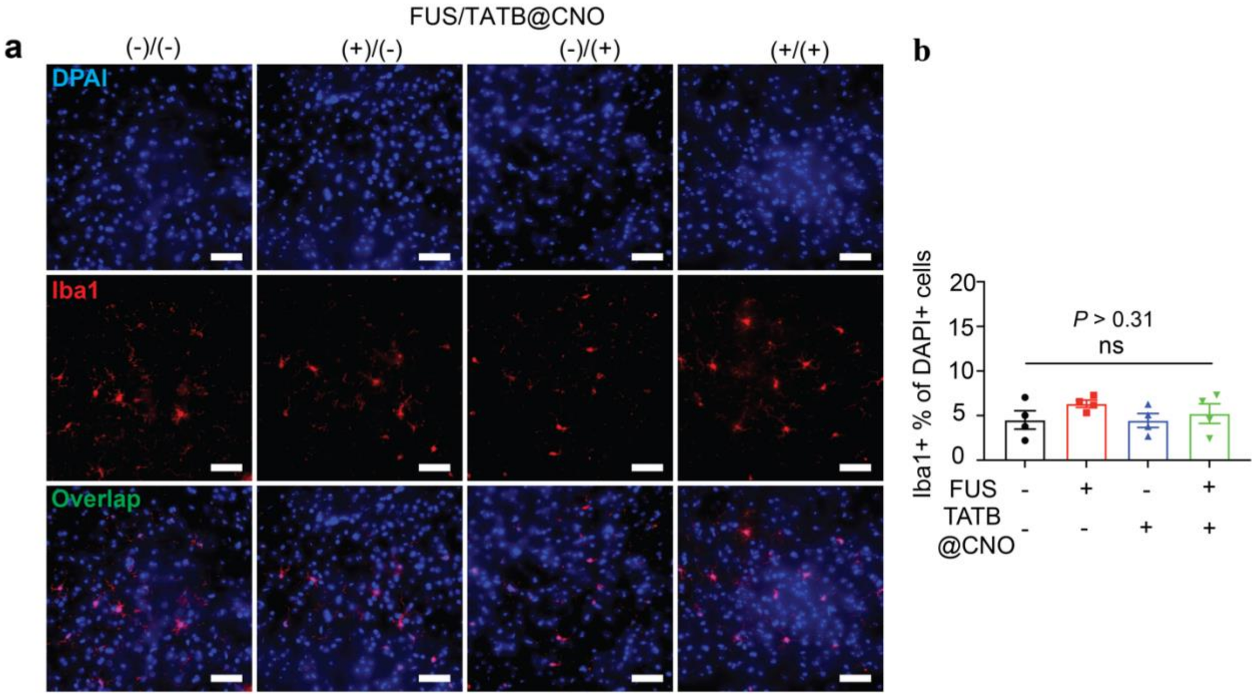


**Supplementary Fig. 24. *In vivo* biocompatibility evaluation of the sono-chemogenetics via determining microglia (Iba1) activation.** (a) Fluorescence images of Iba1 of the brain slices in mouse VTA under the different conditions 7 days after sono-optogenetic stimulation. Scale bar :50 μm. Blue signal is DAPI and red signal is Iba1. (b) Statical analysis of the Iba1 intensity. Mean ± SEM, n >3, mice in each group. Two-way ANOVA and Tukey's multiple comparison test (P ≥ 0.05 (ns), * 0.01≤ P < 0.05, ** 0.001≤ P < 0.01, **** P< 0.0001).

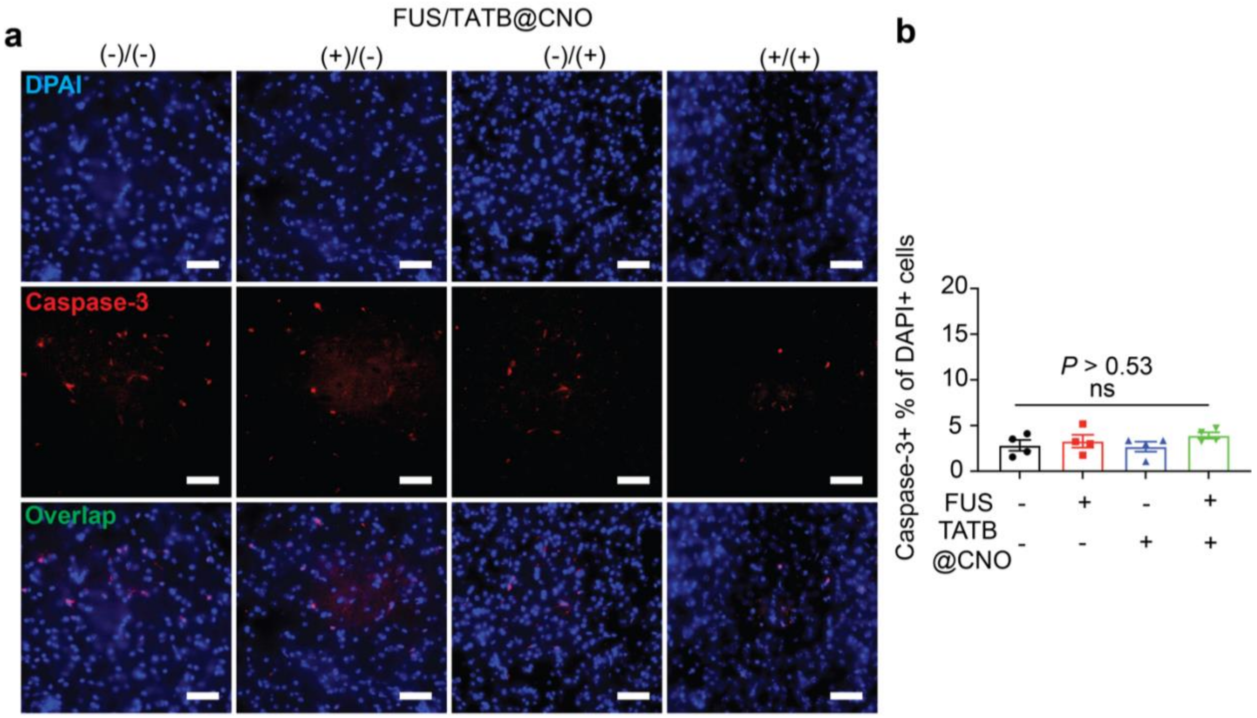


**Supplementary Fig. 25.** ***In vivo*** **biocompatibility evaluation of the sono-chemogenetics via determining neuron apoptosis (Caspase-3).** (a) Fluorescence images of Caspase-3 of the brain slices in mouse VTA under the different conditions 7 days after sono-optogenetic stimulation. Scale bar: 50 μm. Blue signal is DAPI and red signal is Caspase-3. (b) Statical analysis of the Caspase-3 intensity. Mean ± SEM, n >3, mice in each group. Two-way ANOVA and Tukey's multiple comparison test ($P \geq 0.05$ (ns), * $0.01 \leq P < 0.05$, ** $0.001 \leq P < 0.01$, **** $P < 0.0001$).

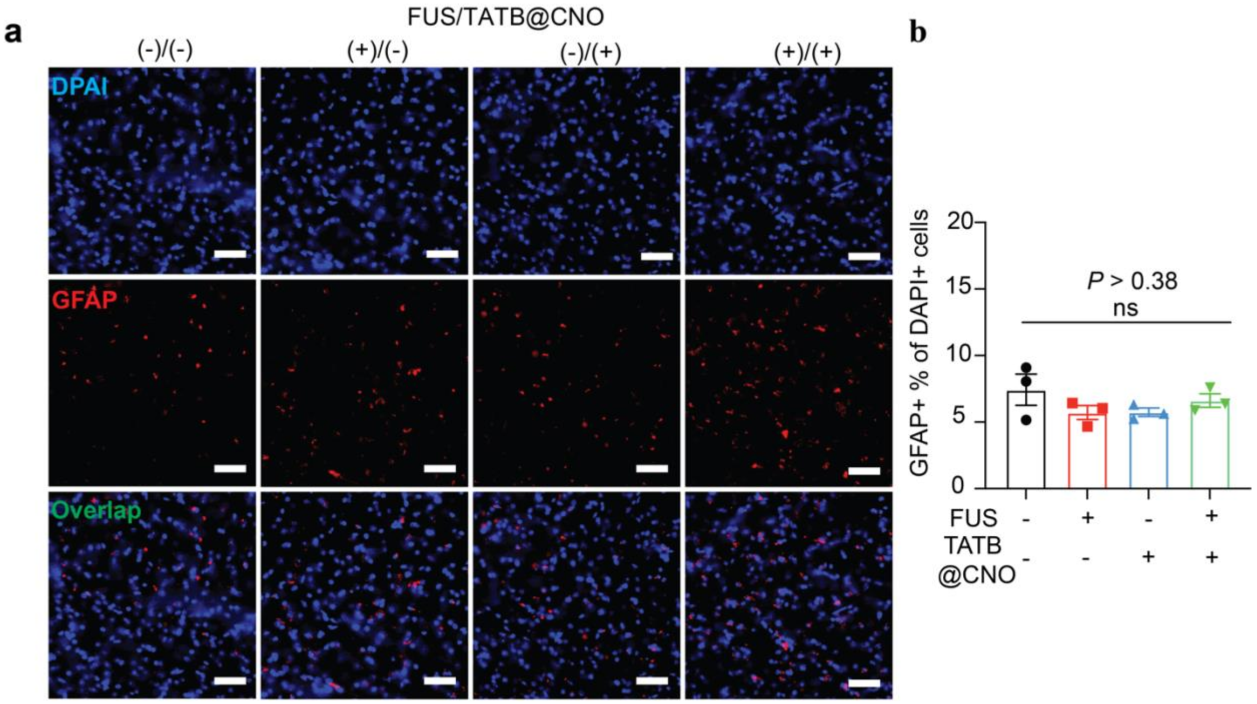


**Supplementary Fig. 26. *In vivo* biocompatibility evaluation of the sono-chemogenetics via determining astrocytes (GFAP) activation.** (a) Fluorescence images of GFAP of the brain slices in mouse VTA under the different conditions 7 days after sono-optogenetic stimulation. Scale bar: 50 µm. Blue signal is DAPI and red signal is GFAP. (b) Statical analysis of the GFAP intensity. Mean ± SEM, n >3, mice in each group. Two-way ANOVA and Tukey's multiple comparison test ($P \geq 0.05$ (ns), * $0.01 \leq P < 0.05$, ** $0.001 \leq P < 0.01$, **** $P < 0.0001$).

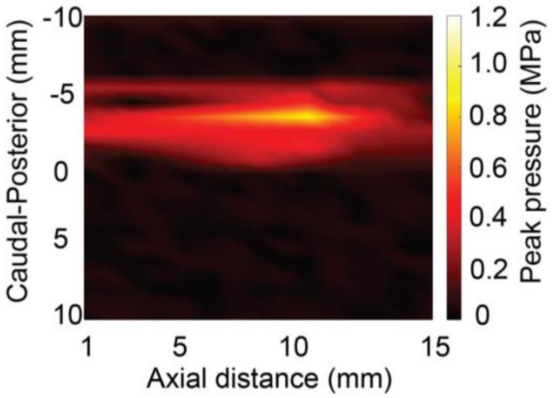


**Supplementary Fig. 27. The *in vivo* ultrasound power transfer in rat heads with FUS focus length of 10 mm.** The ultrasound power heatmap in the rat head shows that around 1.30 MPa was delivered to the rat VTA when 2.45 MPa primary ultrasound power was used.

**Supplementary Table 1**. Summary of reported ultrasound triggered drug delivery systems.

| Materials | Drug loading content (wt%) | Threshold energy for activation. | Ultrasound triggered drug release sensitivity | Percentage of drug release after 20 seconds of stimulation | Stimulation duration required to achieve 20% drug release | Is it possible to programmable modulate ultrasound sensitivity by designing building units? | Reference |
|---|---|---|---|---|---|---|---|
| HOF nanoparticles | 15.1 | 0.51 MPa | Released around 45% after 60 s stimulation at 1.55 MPa | 29 % | ~ 15 s | Yes | This study |
| Lipo–PPIX | < 2.0 | N/A | Released 9% after 10 min stimulation at 3 W $cm^{-2}$ | < 1% | > 30 min | No | *Nat Biomed Eng* **1**, 644–653 (2017) |
| Nanoemulsions | 1.19 | ~ 1.0 MPa | Released around 25%-30% after 60 s stimulation at 1.55 MPa | N/A | N/A | No | *Neuron* **100**, 728–738 (2018) *Nano Lett.* **17**, 652−659 (2017) |
| Disulfide bonded prodrug | N/A | N/A | Release 30%-40% after 60 min stimulation at 15.8 W $cm^{-2}$ | < 1% | > 30 min | No | *Nat. Chem.* **13**, 131–139 (2021) |
| Disulfide-based polymers | N/A | N/A | Release 56% after 180 min stimulation at 15.8 W $cm^{-2}$ | < 1% | > 30 min | No | *Chem. Sci.*, **12**, 1668-1674 (2021) |
| Platinum prodrug | N/A | N/A | Released 10 % after 15 min stimulation at 3.5 W | < 1% | > 30 min | No | *Sci.Adv.*, **9**, eadg5964 (2023) |
| Au-DNA dimer nanoswitches | N/A | N/A | Release 60% after 30 min stimulation at unknow ultrasound power | < 1% | > 5 min | No | *Adv. Sci.* **9**, *2104696 (2022),* |
| Microbubbles | 2.41 | N/A | Immediately releasing 78% under the ultrasound | N/A | N/A | No | *J. Control. Release* **143**, 38–44, (2010) |

**Supplementary Table 2**. Summary of reported non-invasive/minimally-invasive brain stimulation and animal behaviors control.

| Materials | Opsins | Trigger method | Animal species | Maximum depth of brain penetration (mm) | *In vivo* neuron activation in mice | Successful behaviors modulation in mice | *In vivo* neuron activation in rats | Successful behaviors modulation in rats | Reference |
|---|---|---|---|---|---|---|---|---|---|
| HOF nanoparticles | hM3D(Gq) | ultrasound | Mice & rats | 9 | yes | yes | yes | yes | This study |
| / | ChRmine | 635 nm red light | Mice & Rats | 7 | yes | yes | yes | No | *Nat Biotechnol* **39**, 161–164 (2021) |
| magnetic nanoparticles | hM3D(Gq) | magnetic fields | Mice | 5 | yes | yes | No | No | *Nat. Nanotechnol*. **14**, 967–973 (2019) |
| Free CNO | hM3D(Gq) &hM4D (Gi) | ultrasound | Mice | < 5 | yes | yes | No | No | *Nat Biomed Eng* **2**, 475–484 (2018) |
| Propofol loaded Nanoemulsions | N/A | ultrasound | Mice | < 5 | yes | yes | No | No | *Neuron* **100**, 728–738 (2018) *Nano Lett*. **17**, 652−659 (2017) |
| magnetic torquer | Piezo1 | magnetic fields | Mice | < 5 | yes | yes | No | No | *Nat. Mater.* **20**, 1029–1036 (2021). |
| Iron oxide nanoparticles | TRPA1 | magnetic fields | Drosophila melanogaster | < 1 | No | No | No | No | *Nat. Mater.* **21**, 951–958 (2022) |
| / | MscL-G22S | ultrasound | Mice | 5 | yes | yes | No | No | *PNAS* **22**, e2220575120 (2023) |
| / | MscL-G22S | ultrasound | Mice & Rats | < 5 | yes | yes | yes | No | *Nat. Nanotechnol*. **18**, 667–676 (2023) |
| magnetic nanoparticles | TRPV1 | magnetic fields | Mice | 5 | yes | No | No | No | Science, 347, 1477-1480 (2015) |

| | | | | | | | | | |
|---|---|---|---|---|---|---|---|---|---|
| Upconversion nanoparticles | ChR2 | NIR light | Mice | 5 | yes | yes | No | No | *Science* **359**, 679–684 (2018) |
| Phtothermal nanoparticles | TRPV1 | NIR light | Mice | 5 | yes | yes | No | No | *Nat. Biomed. Eng* **6**, 754–770 (2022) |
| Hybrid nanoparticles | ChR2 &KCNQ1 | NIR light | Mice | 5 | yes | yes | No | No | *Adv.Mater.* **35**, 2210018, (2023) |
| Ce:GAGG microparticles | ChR2 | X-ray | Mice | 5 | yes | yes | No | No | *Nat Commun* **12**, 4478 (2021) |
| $Gd_2(WO_4)_3$:Eu nanoparticles | ChR2 | X-ray | Mice | < 5 | yes | No | No | No | *ACS Nano*, **15**, 5201–5208, (2021) |
| / | *hs*TRPA1 | ultrasound | Mice | < 5 | yes | No | No | No | *Nat Commun **13**, 600 (2022)* |
| / | N7T,N308S | ultrasound | Mice | 5 | yes | No | No | No | *Nano Lett.* **20**, 1089–1100, (2020) |
| / | N7T,N308S | ultrasound | Mice | 5 | yes | yes | No | No | *Nano Lett.* **21**, 5967–5976, (2021) |
| Ag/Co-codoped ZnS nanoparticles | ChR2 | ultrasound | Mice | < 5 | yes | yes | No | No | *PNAS* **116**, 26332-26342, (2019) |
| Liposomal nanotransducers | ChR2 | ultrasound | Mice | < 5 | yes | yes | No | No | *J. Am. Chem. Soc* **145**, 1097–1107, (2023) |

**Supplementary Table 3.** Dynamic light scattering tests of HOFs nanoparticles.

| HOFs Nanoparticles | Before ultrasound | | | After ultrasound | | |
|---|---|---|---|---|---|---|
| | Size (nm) | PDI | Zeta potential (mV) | Size (nm) | PDI | Zeta potential (mV) |
| HOF-TATB | 360 ± 6 | 0.35 | -52.8 ± 4.5 | 261 ± 4 | 0.35 | -55.4 ± 2.2 |
| HOF-BTB | 541± 9 | 0.25 | -39.6 ± 1.1 | 358± 1 | 0.21 | -32.6 ± 0.38 |
| HOF-101 | 528± 17 | 0.21 | -34.1 ± 9.8 | 527± 22 | 0.27 | -33.6 ± 10.6 |
| HOF-102 | 306± 11 | 0.31 | -22.7 ± 0.6 | 307± 24 | 0.31 | -44.9± 0.6 |

**Supplementary Table 4.** Structures of relaxed models for HOF-TATB, HOF-BTB, HOF-101 and HOF-102.

| | Dissociated monomer | Hydrogen-bonded dimer | $\pi - \pi$ bonded dimer | HOF |
|---|---|---|---|---|
| HOF-TATB | 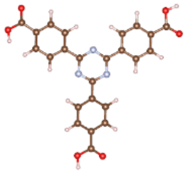 | 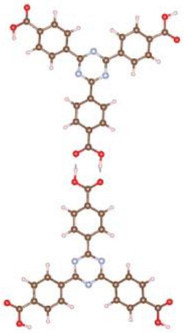 | 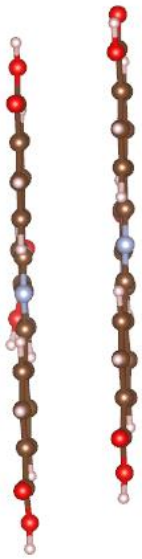 | 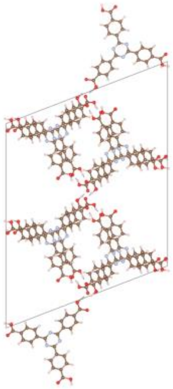 |

| | | | | |
|---|---|---|---|---|
| HOF-BTB | 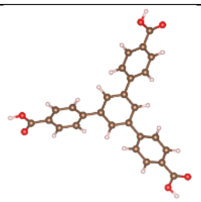 | 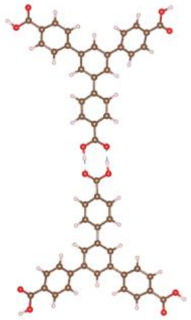 | 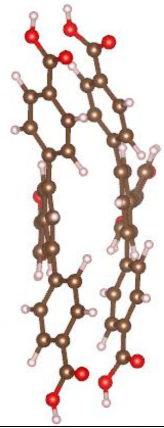 | 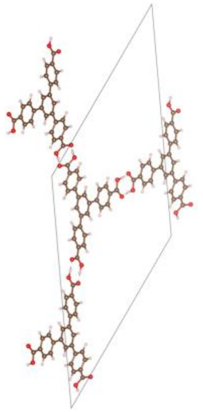 |
| HOF-101 | 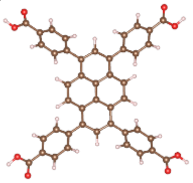 | 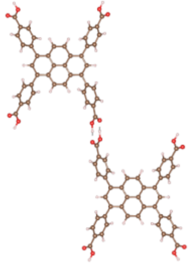 | 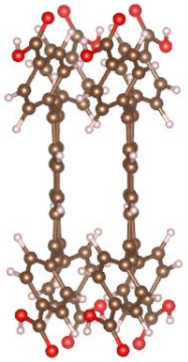 | 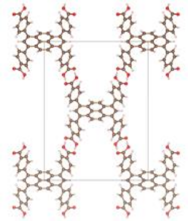 |
| HOF-102 | 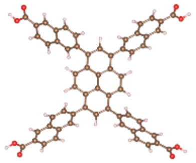 | 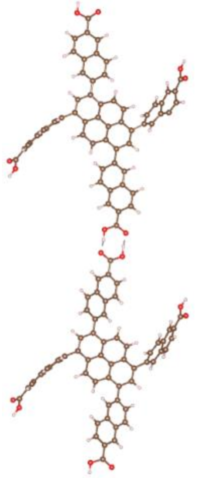 | 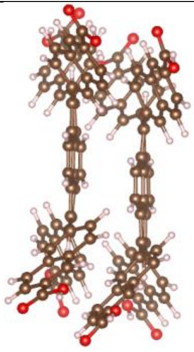 | 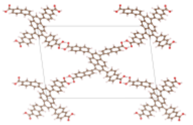 |

**Supplementary Table 5.** The HOFs dissociation constant k at different ultrasound power densities.

| HOFs | Ultrasound peak pressure (MPa) | | | | |
|---|---|---|---|---|---|
| | 1.55 | 1.72 | 3.94 | 6.49 | 8.03 |
| HOF-TATB | 0.2990 | 0.3113 | 0.4194 | 2.0885 | 11.2409 |
| HOF-BTB | 0.1759 | 0.2302 | 0.3996 | 0.4393 | 0.8272 |
| HOF-101 | 0.0057 | 0.0011 | 0.0126 | 0.0611 | 0.0919 |
| HOF-102 | 0.0004 | 0.0009 | 0.0087 | 0.0154 | 0.0492 |

**Supplementary Table 6.** The parameters for the heatmap prediction model.

| Dissociation percentage x | Dissociation Constant k | Ln k | $E_{\underline{US}} = aE_{\text{Cohesive energy}} + b$ | | R squared ($R^2$) |
|---|---|---|---|---|---|
| | | | a | b | |
| 1% | 0.010101 | -4.5951199 | -0.7387 | -4.522 | 0.9529 |
| 2% | 0.020408 | -3.8918203 | -0.7051 | -3.679 | 0.9763 |
| 3% | 0.030928 | -3.4760987 | -0.6853 | -3.181 | 0.9855 |
| 4% | 0.041667 | -3.1780538 | -0.6711 | -2.824 | 0.9892 |
| 5% | 0.052632 | -2.944439 | -0.6599 | -2.544 | 0.9902 |
| 6% | 0.063830 | -2.7515353 | -0.6507 | -2.313 | 0.9896 |
| 7% | 0.075269 | -2.5866893 | -0.6428 | -2.115 | 0.9879 |
| 8% | 0.086957 | -2.442347 | -0.6359 | -1.942 | 0.9855 |
| 9% | 0.098901 | -2.3136349 | -0.6298 | -1.788 | 0.9826 |
| 10% | 0.111111 | -2.1972246 | -0.6242 | -1.648 | 0.9794 |
| 11% | 0.123596 | -2.0907411 | -0.6192 | -1.521 | 0.9759 |
| 12% | 0.136364 | -1.9924302 | -0.6145 | -1.403 | 0.9722 |
| 13% | 0.149425 | -1.9009588 | -0.6101 | -1.293 | 0.9683 |
| 14% | 0.162791 | -1.81529 | -0.606 | -1.191 | 0.9643 |
| 15% | 0.176471 | -1.7346011 | -0.6022 | -1.094 | 0.9602 |

| | | | | | |
|---|---|---|---|---|---|
| 16% | 0.190476 | -1.6582281 | -0.5985 | -1.002 | 0.956 |
| 17% | 0.204819 | -1.5856273 | -0.595 | -0.9153 | 0.9518 |
| 18% | 0.219512 | -1.5163475 | -0.5917 | -0.8322 | 0.9475 |
| 19% | 0.234568 | -1.4500102 | -0.5886 | -0.7527 | 0.9431 |
| 20% | 0.250000 | -1.3862944 | -0.5855 | -0.6764 | 0.9387 |
| 21% | 0.265823 | -1.3249254 | -0.5826 | -0.6028 | 0.9343 |
| 22% | 0.282051 | -1.2656664 | -0.5798 | -0.5318 | 0.9298 |
| 23% | 0.298701 | -1.2083112 | -0.577 | -0.4631 | 0.9253 |
| 24% | 0.315789 | -1.1526795 | -0.5744 | -0.3964 | 0.9207 |
| 25% | 0.333333 | -1.0986123 | -0.5718 | -0.3316 | 0.9162 |
| 26% | 0.351351 | -1.0459686 | -0.5693 | -0.2685 | 0.9116 |
| 27% | 0.369863 | -0.9946226 | -0.5668 | -0.207 | 0.9069 |
| 28% | 0.388889 | -0.9444616 | -0.5644 | -0.1469 | 0.9023 |
| 29% | 0.408451 | -0.895384 | -0.5621 | -0.08804 | 0.8976 |
| 30% | 0.428571 | -0.8472979 | -0.5598 | -0.0304 | 0.8929 |
| 31% | 0.449275 | -0.8001193 | -0.5575 | 0.02613 | 0.8882 |
| 32% | 0.470588 | -0.7537718 | -0.5553 | 0.08168 | 0.8835 |
| 33% | 0.492537 | -0.7081851 | -0.5532 | 0.1363 | 0.8787 |
| 34% | 0.515152 | -0.6632942 | -0.551 | 0.1901 | 0.8739 |

| | | | | | |
|---|---|---|---|---|---|
| 35% | 0.538462 | -0.6190392 | -0.5489 | 0.2432 | 0.8691 |
| 36% | 0.562500 | -0.5753641 | -0.5468 | 0.2955 | 0.8642 |
| 37% | 0.587302 | -0.5322168 | -0.5448 | 0.3472 | 0.8594 |
| 38% | 0.612903 | -0.4895482 | -0.5427 | 0.3984 | 0.8544 |
| 39% | 0.639344 | -0.4473122 | -0.5407 | 0.449 | 0.8495 |
| 40% | 0.666667 | -0.4054651 | -0.5387 | 0.4991 | 0.8445 |
| 41% | 0.694915 | -0.3639654 | -0.5367 | 0.5489 | 0.8395 |
| 42% | 0.724138 | -0.3227734 | -0.5348 | 0.5982 | 0.8344 |
| 43% | 0.754386 | -0.2818512 | -0.5328 | 0.6473 | 0.8293 |
| 44% | 0.785714 | -0.2411621 | -0.5309 | 0.696 | 0.8242 |
| 45% | 0.818182 | -0.2006707 | -0.5289 | 0.7446 | 0.819 |
| 46% | 0.851852 | -0.1603427 | -0.527 | 0.7929 | 0.8137 |
| 47% | 0.886792 | -0.1201443 | -0.5251 | 0.8411 | 0.8085 |
| 48% | 0.923077 | -0.0800427 | -0.5232 | 0.8891 | 0.8031 |
| 49% | 0.960784 | -0.0400053 | -0.5213 | 0.9371 | 0.7977 |
| 50% | 1.000000 | 8.8818E-16 | -0.5193 | 0.9851 | 0.7923 |
| 51% | 1.040816 | 0.04000533 | -0.5174 | 1.033 | 0.7868 |
| 52% | 1.083333 | 0.08004271 | -0.5155 | 1.081 | 0.7812 |
| 53% | 1.127660 | 0.12014431 | -0.5136 | 1.129 | 0.7756 |

| | | | | | |
|---|---|---|---|---|---|
| 54% | 1.173913 | 0.16034265 | -0.5117 | 1.177 | 0.7699 |
| 55% | 1.222222 | 0.2006707 | -0.5098 | 1.226 | 0.7641 |
| 56% | 1.272727 | 0.24116206 | -0.5078 | 1.274 | 0.7583 |
| 57% | 1.325581 | 0.28185115 | -0.5059 | 1.323 | 0.7523 |
| 58% | 1.380952 | 0.32277339 | -0.5039 | 1.372 | 0.7463 |
| 59% | 1.439024 | 0.36396538 | -0.502 | 1.421 | 0.7402 |
| 60% | 1.500000 | 0.40546511 | -0.5 | 1.471 | 0.734 |
| 61% | 1.564103 | 0.44731222 | -0.498 | 1.521 | 0.7277 |
| 62% | 1.631579 | 0.48954823 | -0.496 | 1.572 | 0.7214 |
| 63% | 1.702703 | 0.53221681 | -0.4939 | 1.623 | 0.7148 |
| 64% | 1.777778 | 0.57536414 | -0.4919 | 1.675 | 0.7082 |
| 65% | 1.857143 | 0.61903921 | -0.4898 | 1.727 | 0.7015 |
| 66% | 1.941176 | 0.66329422 | -0.4877 | 1.78 | 0.6946 |
| 67% | 2.030303 | 0.70818506 | -0.4855 | 1.834 | 0.6876 |
| 68% | 2.125000 | 0.7537718 | -0.4834 | 1.888 | 0.6805 |
| 69% | 2.225806 | 0.8001193 | -0.4811 | 1.944 | 0.6731 |
| 70% | 2.333333 | 0.84729786 | -0.4789 | 2.001 | 0.6657 |
| 71% | 2.448276 | 0.89538405 | -0.4766 | 2.058 | 0.658 |
| 72% | 2.571429 | 0.94446161 | -0.4743 | 2.117 | 0.6502 |

| | | | | | |
|---|---|---|---|---|---|
| 73% | 2.703704 | 0.99462258 | -0.4719 | 2.177 | 0.6422 |
| 74% | 2.846154 | 1.04596856 | -0.4694 | 2.239 | 0.6339 |
| 75% | 3.000000 | 1.09861229 | -0.4669 | 2.302 | 0.6254 |
| 76% | 3.166667 | 1.15267951 | -0.4643 | 2.367 | 0.6167 |
| 77% | 3.347826 | 1.20831121 | -0.4617 | 2.433 | 0.6077 |
| 78% | 3.545455 | 1.26566637 | -0.4589 | 2.502 | 0.5984 |
| 79% | 3.761905 | 1.32492541 | -0.4561 | 2.573 | 0.5888 |
| 80% | 4.000000 | 1.38629436 | -0.4532 | 2.647 | 0.5789 |
| 81% | 4.263158 | 1.45001018 | -0.4501 | 2.723 | 0.5686 |
| 82% | 4.555556 | 1.51634749 | -0.447 | 2.802 | 0.5579 |
| 83% | 4.882353 | 1.58562726 | -0.4436 | 2.885 | 0.5467 |
| 84% | 5.250000 | 1.65822808 | -0.4402 | 2.972 | 0.5351 |
| 85% | 5.666667 | 1.73460106 | -0.4365 | 3.064 | 0.5228 |
| 86% | 6.142857 | 1.81528997 | -0.4327 | 3.161 | 0.51 |
| 87% | 6.692308 | 1.90095876 | -0.4286 | 3.263 | 0.4965 |
| 88% | 7.333333 | 1.99243016 | -0.4242 | 3.373 | 0.4821 |

**Supplementary Table 7.** Drug loading content determination.

| HOFs Nanoparticles | Drug loading content (wt%) |
|---|---|
| HOF-TATB | 15.1 ± 1.4 |
| HOF-BTB | 15.8 ± 2.7 |
| HOF-101 | 27.0 ± 1.5 |
| HOF-102 | 29.8 ± 1.3 |

**Supplementary Table 8.** Antibodies used in this work.

| **Primary antibodies** | **Secondary antibodies** |
|---|---|
| Rabbit anti-Iba1<br>(1:500, 013-27691, Wako Chemicals) | Donkey anti-Rabbit, Alexa Fluor 594<br>(1:500, A32754, Invitrogen) |
| Rabbit anti-Cleaved Caspase-3<br>(1:500, 9661, Cell Signaling Tec.) | Donkey anti-Rabbit, Alexa Fluor 594<br>(1:500, A32754, Invitrogen) |
| Rabbit anti-GFAP<br>(1:500, 13-0300, Invitrogen) | Donkey anti-Rabbit, Alexa Fluor 594<br>(1:500, A32754, Invitrogen) |
| Mouse anti-tyrosine hydroxylase antibody (MA1-24654, Fisher Scientific, 1:1000) | Goat anti-mouse Alexa Fluor 488 (1:1000, ab150113, Abcam) |
| Rabbit anti-c-Fos antibody for mice<br>(1:500, ab222699, Abcam) | Goat anti-rabbit Alexa Fluor 405 (1:500, ab175652, Abcam) |
| Rabbit anti-c-Fos antibody for rats (1:500, ab289723, Abcam) | Goat anti-rabbit Alexa Fluor 405 (1:500, ab175652, Abcam) |
| | H&E staining kit<br>(ab245880, Abcam) |
| | Hoechst 33342<br>(1:5000, 17535, AAT Bioquest ) |

**References:**


1. Chen, T.-W. *et al.* Ultrasensitive fluorescent proteins for imaging neuronal activity. *Nature* **499**, 295–300 (2013).
